# The intestinal circadian clock drives microbial rhythmicity to maintain gastrointestinal homeostasis

**DOI:** 10.1101/2021.10.18.464061

**Authors:** Marjolein Heddes, Baraa Altaha, Yunhui Niu, Sandra Reitmeier, Karin Kleigrewe, Dirk Haller, Silke Kiessling

## Abstract

Diurnal (*i.e*., 24-hour) oscillations of the gut microbiome have been described in various species including mice and humans. However, the driving force behind these rhythms remains less clear. In this study, we differentiate between endogenous and exogenous time cues driving microbial rhythms. Our results demonstrate that fecal microbial oscillations are maintained in mice kept in the absence of light, supporting a role of the host’s circadian system rather than representing a diurnal response to environmental changes. Intestinal epithelial cell-specific ablation of the core clock gene *Bmal1* disrupts rhythmicity of microbiota. Targeted metabolomics functionally link intestinal clock-controlled bacteria to microbial-derived products, in particular branched-chain fatty acids and secondary bile acids. Microbiota transfer from intestinal clock-deficient mice into germ-free mice altered intestinal gene expression, enhanced lymphoid organ weights and suppressed immune cell recruitment. These results highlight the importance of functional intestinal clocks for circadian microbiota composition and function, which is required to balance the host’s gastrointestinal homeostasis.

## Introduction

Various physiological processes show 24-hour fluctuations. These rhythms are the expression of endogenous circadian (Lat. circa = about; dies = day) clocks that have evolved in most species ^1^ to facilitate anticipation of daily recurring environmental changes. In mammals, the circadian system includes a central pacemaker regulating sleep-wake behavior and orchestrating subordinated tissue clocks by humoral and neuronal pathways ^2^. At the cellular level, these clocks consist of inter-regulated core clock genes ^3^ that drive tissue-specific transcriptional programs of clock-controlled genes (CCGs) ^4^. Through these CCGs, circadian clocks regulate various aspects of physiology including metabolism, gastrointestinal transit time (GITT), mucus secretion, antimicrobial peptide secretion, immune defense and intestinal barrier function (reviewed by ^5–8^).

In mice, 10-15 % of gut bacteria undergo diurnal oscillations in their abundance influenced by meal timing, diet type and other environmental conditions ^9–11^. Recently, we found similar rhythms in microbiota composition and function in population-based human cohorts ^12^. Although functionality of the host’s circadian system impacts microbial rhythmicity ^10,13,14^, it is unclear which tissue clocks contribute to this effect.

A balanced gut microbiome promotes health and microbial dysbiosis has been linked to metabolic diseases, colorectal cancer and gastrointestinal inflammation ^12,15–17^. Similar pathological consequences are associated with circadian rhythm disruption (reviewed by ^18–20^), which also induces microbiota dysbiosis ^10,11,13,14,21^. Consequently, we hypothesized that circadian regulation of microbiota composition and function may contribute to the host’s GI health.

Here, we functionally dissect the circadian origin of microbiota oscillations in mice. We provide evidence that intestinal epithelial cell (IEC) clocks generate the majority of gut microbial rhythms and their metabolic products, particularly short-chain fatty acids (SCFAs) and bile acids (BAs). Transfer of microbiota from IEC clock-deficient mice in germ-free (GF) wild type hosts directly indicate the consequences of microbiota arrhythmicity on the gastrointestinal homeostasis. Thus, we identify a mechanistic link between IEC clocks, gut bacteria rhythms and their functions through transfer experiments, showing the importance of rhythmic microbiota for host physiology.

## Results

### Rhythms in microbiota are generated endogenously by the circadian system

Diurnal rhythms in microbiota composition and function have been demonstrated in animal models and in humans ^9–12^. However, it has not been demonstrated whether these rhythms are a response to rhythmic external cues (*Zeitgebers*), such as the light-dark cycle, or are generated by endogenous clocks ^22^ and, thus, persist when the organism is placed in an environment devoid of timing cues. To address this question, we compared fecal microbiota rhythms of the same wild-type mice kept in a rhythmic 12-hour light/12-hour dark (LD) cycle and constant darkness (DD) for two weeks (**Fig. 1A**). Host-driven rhythmic factors, which might influence microbiota composition, such as locomotor activity, food intake as well as total GI transit time (GITT), did not differ between light conditions (**Suppl. Fig 1A-D**). 16S rRNA profiling of fecal samples revealed clustering based on sampling time points rather than light condition (**Fig. 1B, 1E**). Generalized UniFrac distances (GUniFrac) quantification to ZT1 identified rhythmicity in both light conditions (**Fig. 1C**). Importantly, 24-hour rhythms of species richness and Shannon effective number of species found in LD persisted in DD, supporting their circadian origin (**Fig. 1D**). Rhythms were also preserved in DD on phylum level with the two most dominant phyla Bacteroidetes and Firmicutes, oscillating in antiphase (**Fig. 1E, F)**. Although microbiota composition is commonly analyzed by relative abundance, rhythmicity of highly abundant taxa may mask oscillations of small microbial communities, as previously demonstrated in fecal samples collected in LD ^13^. Therefore, using synthetic DNA spikes, we performed relative quantification of the copy number of 16S rRNA genes according to Tourlousee et al. ^23^, from here on referred to as ‘quantitative analysis’. With both approaches, highly abundant phyla and families, including *Lachnospiraceae* and *Muribaculaceae*, showed comparable circadian rhythmicity in LD and DD (**Fig. 1F; Suppl. Fig. 1E**). Few families, such as *Prevotellaceae* showed significant diurnal (LD), but no circadian (DD) rhythmicity, suggesting that their rhythms are regulated by the environmental LD cycle (**Suppl. Fig. 1E**). After removal of low-abundant taxa (mean relative abundance > 0.1%; prevalence > 10%), the remaining 580 zero-radius OTUs (zOTUs) displayed robust and comparable 24-hour oscillations in both light conditions, independent of the analysis (**Fig. 1G, H; Suppl. Fig. 1F-I**). The wide distribution of peak abundances, e.g., bacteria peaking during the day (mainly belonging to *Muribaculaceae*) and during the night (*Lachnospiraceae*) (**Fig. 1G, H; Suppl. Fig. 1F-H**), suggests that different microbial taxa dominate different daytimes. Importantly, more than 60% of all identified zOTUs were found to significantly oscillate in LD and in DD (**Fig. 1G, H, Suppl. Fig. 1F-H, Suppl. table 1-3**), suggesting that their rhythmicity is generated by the circadian system. Notably, comparable amounts of rhythmic zOTUs belong to the major phyla Firmicutes and Bacteriodetes (**Suppl. Table 2**). Previous literature identified lower percentages of microbial rhythmicity ^9,10^. To assess this discrepancy we compared our results with the previously published data set from Thaiss and colleagues ^24^. Although the different methods to assign taxonomic units, namely OTUs (97 % similarity strain assignment) in contrast to zOTUS (100% similarity strain assignment) caused disparity between amounts of rhythmic OTUs/zOTUs, the total abundance of rhythmic microbiota was comparable between both studies (Heddes et al: 74 %, Thaiss et al: 83%, **Suppl. Fig. 1L. Suppl. Table 4**). Rhythmicity of the majority of zOTUs (e.g. the genera *Alistipes, Oscillibacter* and *Fusimonas*) identified in our study by relative analysis was further validated by quantitative analysis (73 % in LD, 58 % in DD) **(Suppl. Fig. 1 I, J).** The diurnal rhythmicity found in ¾ of all zOTUs examined in LD using JTK_CYCLE ^25^ is generated by the endogenous circadian system, since DODR analysis ^26 27^confirmed that more than 95% of zOTUs show similar rhythmicity in DD (e.g. the genera *Eubacterium* and members of the family *Lachnospiraceae*) (**Fig. 1G**; **Suppl. Fig. 1K, Suppl. Table 1, Suppl. Table 3)**.

**Figure 1.**
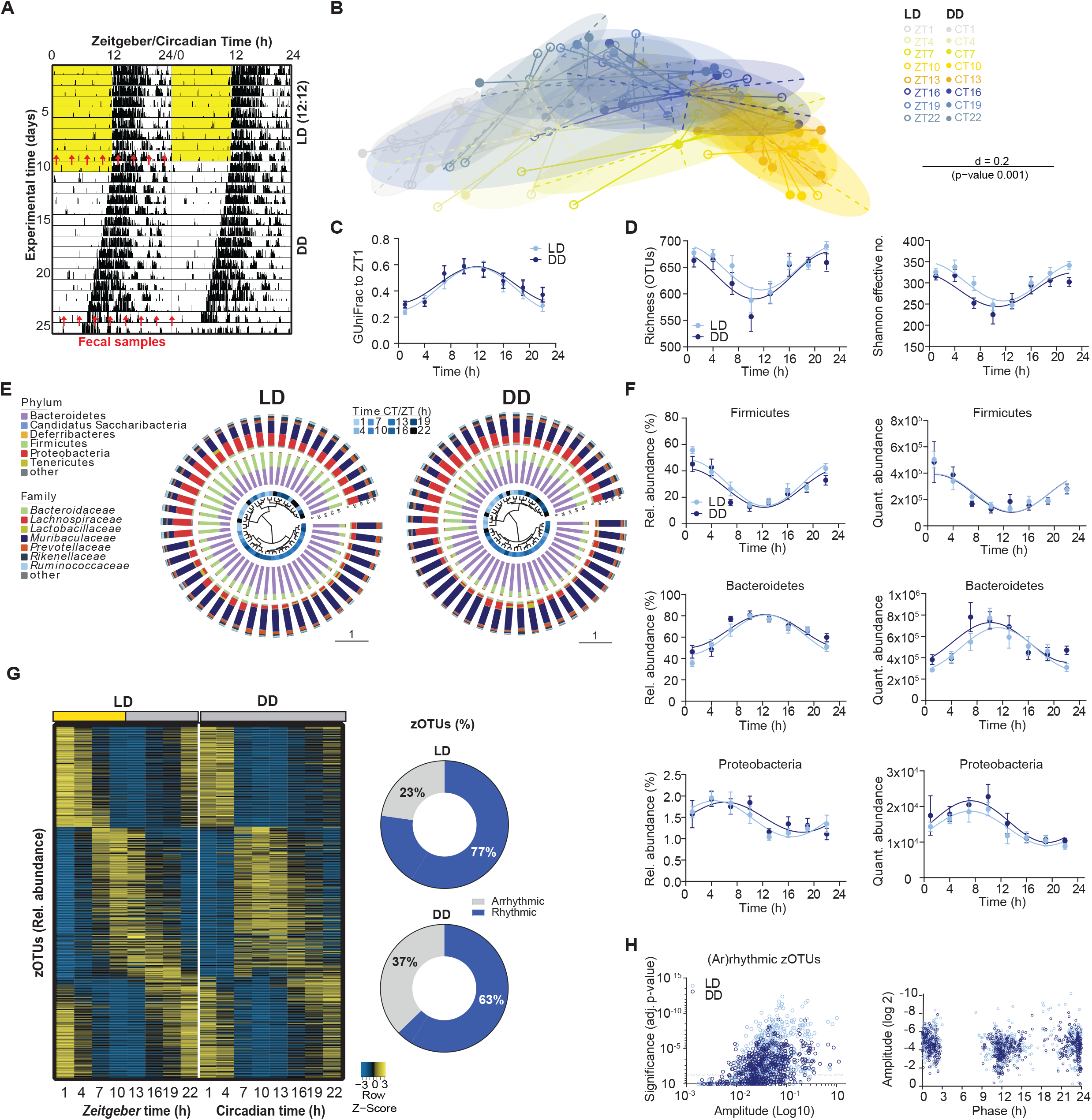
Fecal microbial rhythms persist in constant darkness. (A) Representative actogram of control (*Bmal1^IECfl/fl^*) mice exposed to light-dark (LD) cycle and two weeks of constant darkness (DD). Fecal sampling times are indicated by red arrows. (B) Beta-diversity principal coordinates analyses plot (PCoA) of fecal microbiota based on generalized UniFrac distances (GUniFrac) in LD and DD conditions and its quantification over time relative to ZT1 (C). (D) Diurnal (LD) and circadian (DD) profiles of alpha-diversity. (E) Dendrogram of microbiota profiles based on GUniFrac in LD and DD conditions. Taxonomic composition at phylum and family level for each sample is shown as stacked bar plots around the dendrogram. The blue inner circle indicates the sampling time. (F) Diurnal (LD) and circadian (DD) profiles of the relative (left) and quantitative abundance (right) of phyla. (G) Heatmap depicting the relative abundance of 580 zOTUs (mean relative abundance > 0.1%; prevalence > 10%). Data are normalized to the peak of each zOTU and ordered by the peak phase in LD conditions. Pie-charts at the right indicate the amount of rhythmic (blue) and arrhythmic (grey) zOTUs identified by JTK_Cycle (adj. p-value ≤ 0.05) (H) Significance and amplitude of rhythmic and arrhythmic zOTUs (left) and phase distribution (right) in LD and DD based on relative analysis. Significant rhythms are illustrated with fitted cosine-wave regression using a solid line (significance: p-value ≤ 0.05. LD (light-blue) and DD (dark-blue). n = 6 mice/time point/light condition. Data are represented as mean ± SEM.

### Robust microbial rhythms despite the absence of rhythmic food intake behavior

Light and food are the main *Zeitgeber* for the central and peripheral clocks, respectively ^2,28^. In our study, external light conditions only mildly effected microbial rhythms (**Fig. 1**) and previous research showed that rhythmic feeding time influences the phasing of microbiota rhythmicity ^10,11^. This prompted us to determine the dependency of bacterial oscillation on rhythmic food intake behavior by investigating microbiota rhythmicity in the absence of both *Zeitgeber* food and light. Therefore we collected feces from starved mice during the 2^nd^ day in DD (**Fig. 2A**). Although starvation altered microbial composition, clustering according to collection time points was significant in both feeding conditions (**Fig. 2B, D**). Microbiota diversity became arrhythmic in the absence of food (**Fig. 2C)**. However, rhythms persisted in the two dominant phyla, highly abundant families, including *Muribaculaceae* and *Lachnospiraceae* and the majority of zOTUs (**Fig. 2E-H, Suppl. Table 1)**. Of note, circadian food intake behavior enhanced the amount of rhythmic zOTUs by 16% according to DODR analysis on quantitative data (**Suppl. Table 3**). Nevertheless, circadian rhythms of highly abundant microbial taxa belonging to *Muribaculaceae* and *Ruminococaceae* maintained under starving conditions in both relative and quantitative analysis (**Fig. 2 E-I)**.

**Figure 2.**
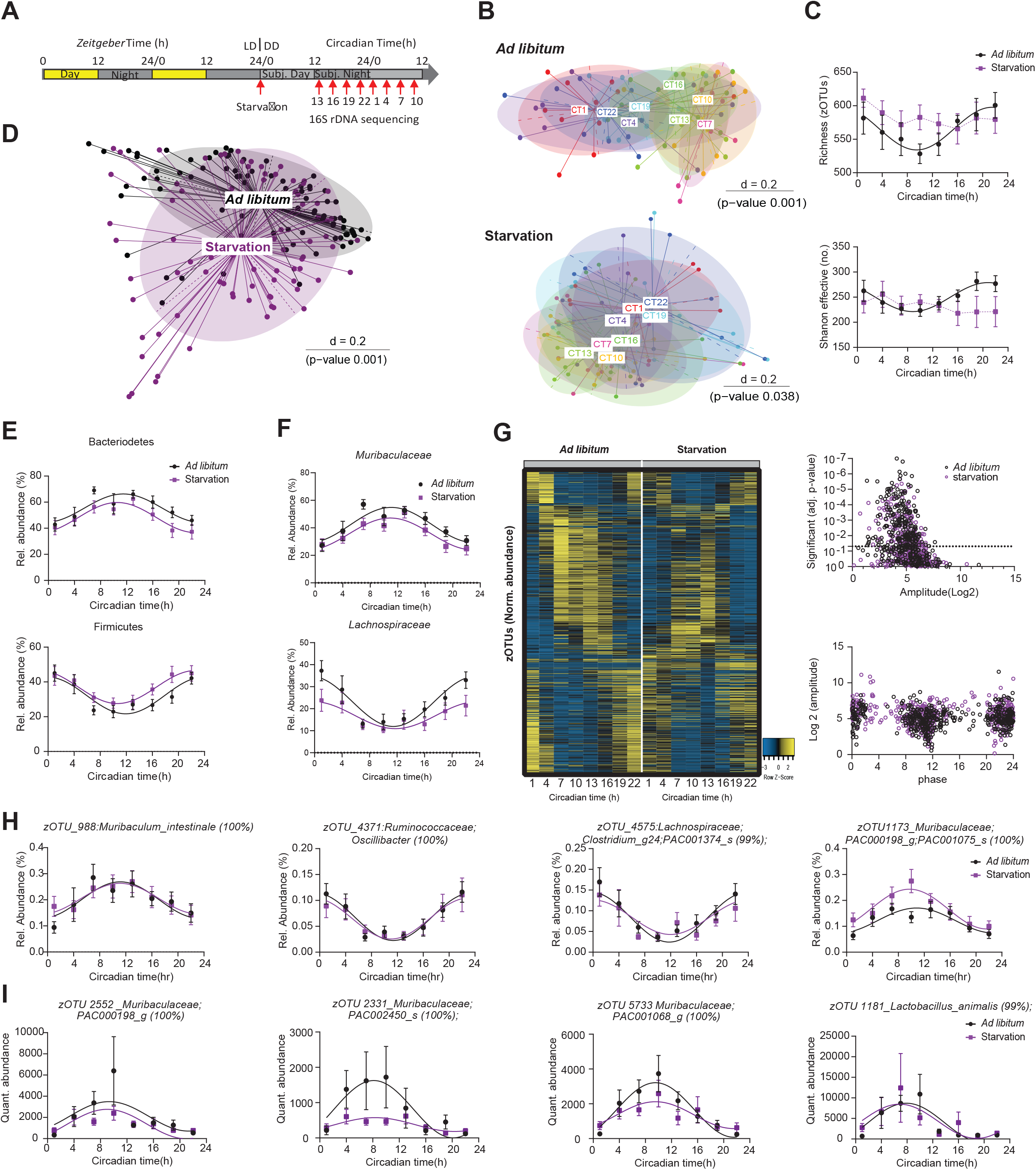
Food intake behavior marginally masks microbiota rhythmicity. (A) Schematic illustration of the experiment design. (B) Beta-diversity principal coordinates analyses plots (PCoA) on generalized UniFrac distances (GUniFrac) distances of fecal microbiota stratified by individual time points in *ad libitum* (top) and starvation (buttom). (C) Circadian profiles of alpha-diversity. (D) PCoA plots of fecal microbiota based on GUniFrac distances stratified by feeding condition. (E, F) Circadian profiles of the relative abundance of the major phyla (E) and family (F) of fecal microbiota. (G) Heatmap depicting the relative abundance of 511 zOTUs (mean relative abundance > 0.1%; prevalence > 10%). Data are normalized to the peak of each zOTU and ordered by the peak phase in ad libitum condition. Significance and amplitude (JTK_Cycle analysis) of rhythmic and arrhythmic zOTUs (top) and phase distribution (based on cosine regression analysis) (bottom). Dashed line indicates adj p-value = 0.05. (H, I) Circadian profile of relative (H) and quantitative (I) abundance of zOTUs of fecal microbiota. Significant rhythms (cosine-wave regression, p-value ≤ 0.05) are illustrated with fitted cosine-wave curves; data points connected by dotted lines indicate no significant cosine fit curves (p-value > 0.05) and thus no rhythmicity. zOTUs were further analyzed with the adjusted compare rhythm script based on DODR. n = 9-11 mice/time point/light condition for relative data. n = 4-5 mice/time point/light condition for quantitative data. Data are represented by mean ± SEM.

Together, these data show that although rhythms in gut bacteria are slightly influenced by environmental light and food conditions, intrinsic factors are the dominant drivers of circadian microbiota fluctuations.

### The intestinal circadian clock controls microbiota composition and function

An important role of the host clock on diurnal microbiota fluctuation has previously been suggested based on experiments on mice with clock dysfunction in all tissues under LD conditions ^10,13,14^. The GI tract represents an important interface for cross-talk between IECs, bacteria and their metabolites ^29^ and thus might likely be involved in controlling microbiota composition. To identify the circadian origin of microbial rhythmicity, we used mice with IEC-specific deletion of the essential core clock gene *Bmal1* (*Bmal1^IEC-/-^*). The rhythmic activity, food intake behavior as well as GITT and stool weight of *Bmal1^IEC-/-^* mice did not differ from control littermates, reflecting a functional central pacemaker (**Suppl. Fig. 2A-E)**. Arrhythmic core clock gene expression in the jejunum, cecum and proximal colon, in contrast to rhythms in the liver, confirmed intestine-specific clock ablation in *Bmal1^IEC-/-^* mice (**Suppl. Fig. 2F**). Rhythmic genes relevant for host-microbe crosstalk, in particular *Tlr2*, *Muc2*, *Nfil3* and *Hdac3*^30–35^ lost rhythmicity in *Bmal1^IEC-/-^* mice (**Suppl. Fig. 2G**). In line with this, microbiota composition in DD and LD differed significantly between *Bmal1^IEC-/-^* mice and controls (p = 0.037 DD, p=0.018 LD) and circadian rhythmicity in community diversity (species richness) observed in control mice was abolished in *Bmal1^IEC-/-^* mice (**Fig. 3A, B, Fig. 4A**). Quantitative analysis revealed loss of rhythmicity of Firmicutes in *Bmal1^IEC-/-^* mice in LD and DD (LD p=, DD p = 0.313), although, oscillation in relative abundance of the major phyla was found in both genotypes, (**Fig. 3C, Fig. 4C**). Alterations in quantitative compared to relative rhythmicity analyses are not due to differences in 16s copy number over time, as both genotypes show similar patterns in LD and DD conditions (**Suppl. Fig. 2H, Fig. 4B**). In accordance to results obtained from whole-body clock-deficient mice ^13^, IEC-specific loss of *Bmal1* decreased the quantitative abundance of both phyla, which was not visible in relative data sets (**Fig. 3C, Fig. 4C**). Similarly, rhythmicity observed in the relative abundance of families, including *Lachnospiraceae* and *Ruminococcaceae*, was abolished performing quantitative analysis (**Suppl. Fig. 2I).** Furthermore, dramatically disrupted circadian oscillations of *Bmal1^IEC-/-^* mice in both relative and quantitative analyses was observed on the level of all 580 identified zOTUs illustrated by heatmaps (**Fig. 3D, E**). Of note, in *Bmal1^IEC-/-^* mice the phase of remaining microbial oscillations was delayed using quantitative analysis (**Fig. 3D, E)**. Based on both analyses more than 60 % of fecal zOTUs, which account for more than two-third of bacterial abundance, underwent circadian oscillation in controls (**Fig. 3D, E, Suppl. Table 1**). Importantly, the amount of rhythmic zOTUs was dramatically reduced by ⅔ in mice with IEC-specific *Bmal1* deficiency in both analyses, independent of the light conditions (**Fig. 3D, E, Fig. 4D, I, Suppl. Table 1-3**. Circadian zOTUs in control mice, which were arrhythmic in *Bmal1^IEC-/-^* mice in both relative and quantitative analysis, belong predominantly to the families *Lachnospiraceae* and *Ruminococcaceae* of the phylum Firmicutes (**Suppl. Fig. 3J, Suppl. Table 2**). The genera *Lactobacillus, Ruminococcus, Anaerotignum, Odoribacter* and *Alistipes* are among the taxa with differential rhythmicity comparing *Bmal1^IEC-/-^* mice with their controls based on DODR analysis and in addition significantly differed in their abundance between genotypes (**Fig. 3F, G**). Although the majority of zOTUs was under intestinal clock control, rhythmicity of around 20% of zOTUs (e.g. *Lactobacillus animalis* and *Pseudoflavonifractor*)persisted in *Bmal1^IEC-/-^* mice (**Fig. 3D, E, Suppl. Table 1**). To address whether these remaining rhythms were driven by the rhythmic food intake behavior observed in *Bmal1^IEC-/-^* mice kept under ad libitum conditions in DD (**Suppl. Fig. 2D**), microbiota rhythmicity in *Bmal1^IEC-/-^* mice was analyzed in feces from starved mice during the 2^nd^ day in DD (**Fig. 2A**). Although beta diversity was significantly altered in the absence of rhythmic food intake profiles of major phyla were undistinguishable between feeding conditions **(Fig. 4E, F).** Only a small amount of the remaining rhythmic zOTUs in *Bmal1^IEC-/-^* mice significantly lost rhythmicity upon starvation based on DODR analysis **(Suppl. Table 3)**, such as bacteria belonging to *Turicimonas muris* and *Muribaculaceae*, and (**Fig. 4G-I, Suppl. Table 1**).

**Figure 3.**
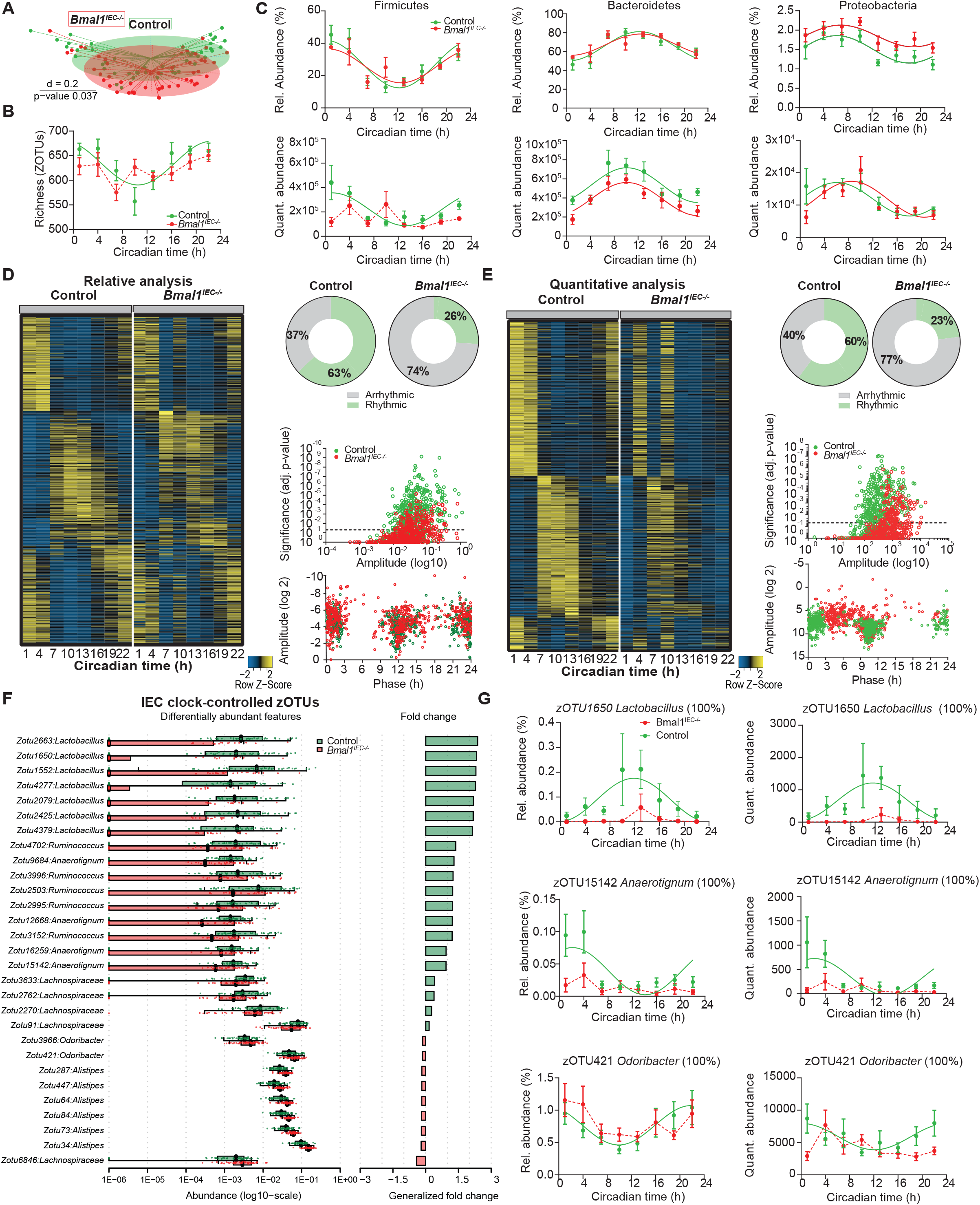
The intestinal circadian clock drives circadian microbiota composition. (A) Beta-diversity principal coordinates analyses plot (PCoA) of fecal microbiota based on generalized UniFrac distances (GUniFrac) stratified by genotype. (B) Circadian profile of alpha diversity. (C) Circadian profile of relative (top) and quantitative (bottom) abundance of the major phyla. (D, E) Heatmap depicting the relative abundance (D) and quantitative abundance (E) of 580 zOTUs (mean relative abundance > 0.1%; prevalence >10%). Data are normalized to the peak of each zOTU and ordered by the peak phase of control mice. Pie-charts at the right indicate the amount of rhythmic (colored) and arrhythmic (grey) zOTUs identified by JTK_Cycle (rhythmic = adj. p-value ≤ 0.05) based on relative (D) and quantitative (E) analysis. Significance and amplitude of rhythmic and arrhythmic zOTUs (top) and phase distribution (bottom) in controls and intestinal epithelial specific Bmal1-deficient (*Bmal1^IEC-/-^*) mice is depicted on the right of the heatmaps. Dashed line indicates adj. p-value = 0.05 (JTK_Cycle). Bar charts in (F) represent the intestinal controlled zOTUs (adjusted compare rhythm script based on DODR) abundance comparison between control and *Bmal1^IEC-/-^*(*Wilcoxon*, adj. p-value ≤ 0.05). Box and bar plots illustrate the alteration in quant. abundance (adj. p-value ≤ 0.05) and fold change of gut controlled zOTUs in the fecal samples with examples depicted in (G). Significant rhythms are illustrated with fitted cosine-regression solid lines; data points connected by dotted lines indicate no significant cosine fit curves (p-value > 0.05) and thus no rhythmicity. *Bmal1^IECfl/fl^* controls (green) and *Bmal1^IEC-/-^* (red). n = 5-6 mice/time point/genotype. Data are represented as mean ± SEM.

**Figure 4.**
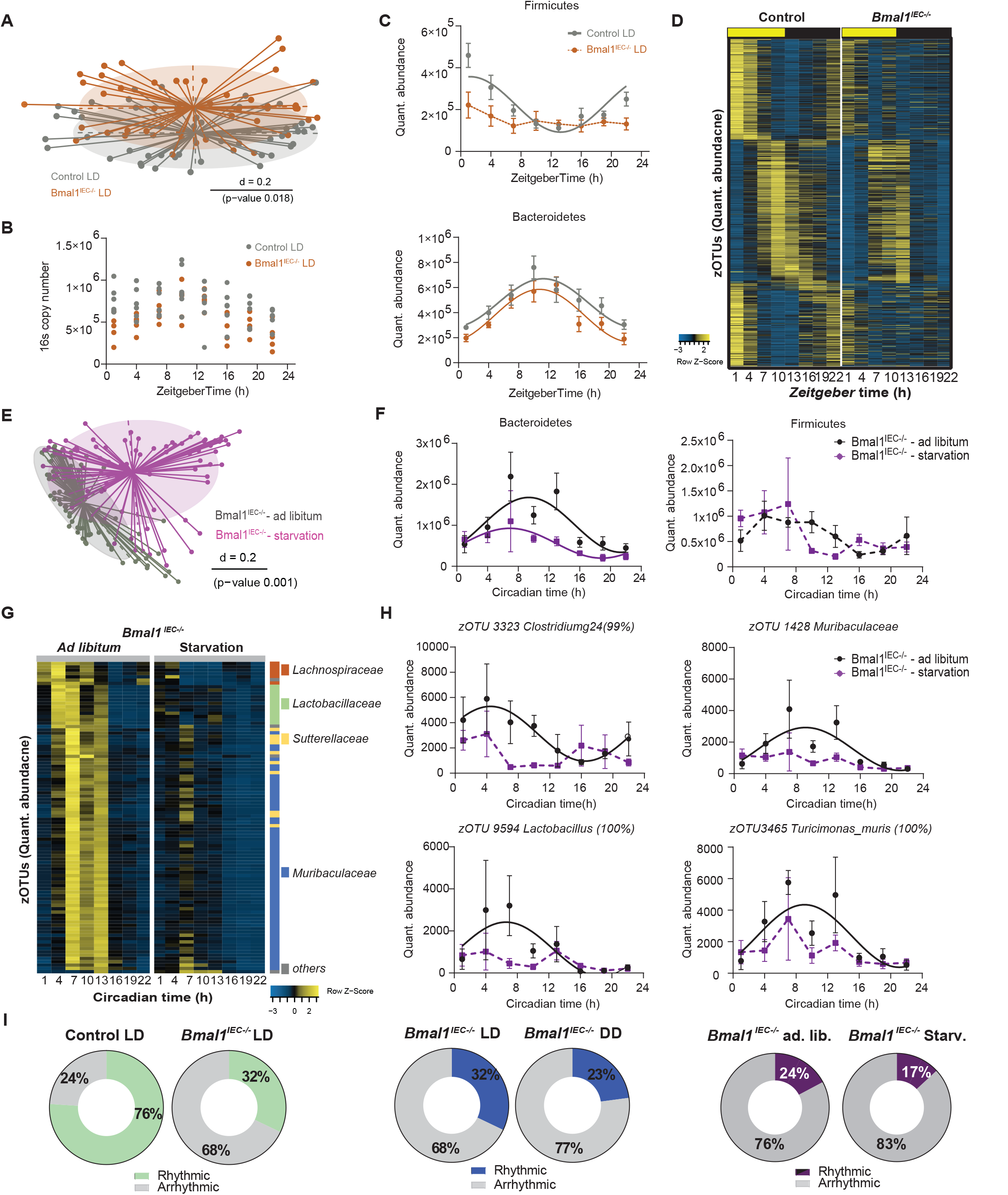
Microbiota profiling in *Bmal1^IEC-/-^* mice in LD conditions and under starvation. (A) Beta-diversity principal coordinates analyses plot (PCoA) based on generalized UniFrac distances (GUniFrac) of fecal microbiota of *Bmal1^IEC-/-^* and their *Bmal1^IECfl/fl^* controls in LD conditons as well as the diurnal profiles of (B) 16s copy number (2-way ANOVA) and (C) major phyla quantitative abundance. (D) Heatmap illustrating zOTUs quantitative abundance over time of fecal microbiota of *Bmal1^IEC-/-^* and their controls in LD conditions. Data are ordered by the peak phase in LD conditions of the control group. (E) Beta-diversity principal coordinates analyses plot (PCoA) based on generalized UniFrac distances (GUniFrac) of fecal microbiota stratified by food availability. (F) Circadian profiles of phyla Firmicutes and Bacteroidetes. (G) Heatmap illustrating microbiota loosing/changing rhythmicity after starvation in *Bmal1^IEC-/-^* mice according to the adjusted compare rhythm script based on DODR (adj. p-value ≤ 0.05). Data are normalized to the peak of each zOTU and ordered by the peak phase in ad libitum condition. (H) Circadian profiles of the quantitative abundance of example zOTUs masked by the food-intake behavior (adjusted compare rhythm script based on DODR, adj. p-value ≤ 0.05). (I) Pie charts indicating the percentage of rhythmic/arrhythmic zOTUs in different condition according to JTK cycle analysis (rhythmic = adj. p-value ≤ 0.05). Significant rhythms (Cosine regression p-value ≤0.05) are illustrated with fitted cosine-regression; data points connected by dotted lines indicate no significant cosine fit curves (p-value > 0.05) and thus no rhythmicity. zOTUs were further analyzed with the adjusted compare rhythm script based on DODR. n = 3-6 mice/time point/condition. Data are represented as mean ± SEM

Taken together, rhythmicity analyses of zOTU in *Bmal1^IEC-/-^* mice and controls under different light and food conditions (**Fig. 3, Fig. 4, Suppl. Fig. 2**) highlight that the IEC clock represent the dominant driver of circadian gut microbiota oscillation.

### Intestinal clock-controlled microbial functions balance gastrointestinal homeostasis

To address the potential physiological relevance of microbial oscillations, PICRUST 2.0 analysis was performed on intestinal-clock controlled zOTUs. Loss of microbial rhythmicity in *Bmal1^IEC-/-^* mice was reflected in assigned pathways involved in sugar- and amino acid metabolism, vitamin biosynthesis and relevant for fatty acid (FA) metabolism such as β-oxidation, FA elongation and short-chain FA (SCFA) fermentation (**Fig 5A, Suppl. Fig 3 I**). This prompted us to test the functional connection between intestinal clock-driven microbial rhythmicity and intestinal homeostasis (**Fig. 5B-H**). Indeed, Procrustes analyses (PA) identified an association between intestinal clock-controlled zOTUs and SCFA concentrations measured by targeted metabolomics (p = 0.001) (**Fig. 5B**). The level of valeric acid (p = 0.04, Mann-Whitney U test) and low-abundant branched-chain fatty acids (BCFAs) (p = 0.01, Mann-Whitney U test), including isovaleric, isobutyric, 2-methylbutyric differed between genotypes, although the amount of total and highly abundant SCFAs were comparable (**Fig 5C, Suppl. Fig. 3A**). Nevertheless, concentrations of SCFAs negatively correlated with the relative abundance of intestinal clock-driven taxa, mainly belonging to the family *Muribaculaceae* of the phylum Bacteroidetes (**Fig. 5D, Suppl. Fig. 3B**). Positive significant correlations with multiple SCFAs, in particular BCFAs, where found with zOTUs belonging to the phylum Firmicutes, including the SCFA producers *Lachnospiraceae* and *Ruminococcaceae* (**Suppl. Fig. 3B**) ^36^. Similar to the results observed for SCFAs, associations were found between intestinal clock-controlled zOTUs and BAs levels (p = 0.04, **Fig. 5G**). Positive correlations with primary BAs and negative correlations with secondary BAs were observed with several intestinal clock-controlled taxonomic members mainly belonging to Firmicutes (**Fig. 5D, Suppl. Fig. 3D**). For example, DHLCA and 7-sulfo-CA negatively correlated with *Oscillibacter* and *Eubacterium* (**Fig. 5D, Suppl. Fig. 3D**). In addition, almost half of the measured BAs differed among genotypes and, BAs, such as the conjugated primary BA, TCDA, and secondary BA, 7-sulfo-CA, lost rhythmicity in *Bmal1^IEC-/-^* mice (**Fig. 5E-F**). Interestingly, most alterations were observed for secondary BAs (**Fig. 5H, Suppl. Fig3 C**), which were linked to various GI diseases (reviewed by ^37^). In summary, these results indicate that deletion of intestinal clock function causes loss of rhythmicity in the majority of bacterial taxa and alters microbial functionality.

**Figure 5.**
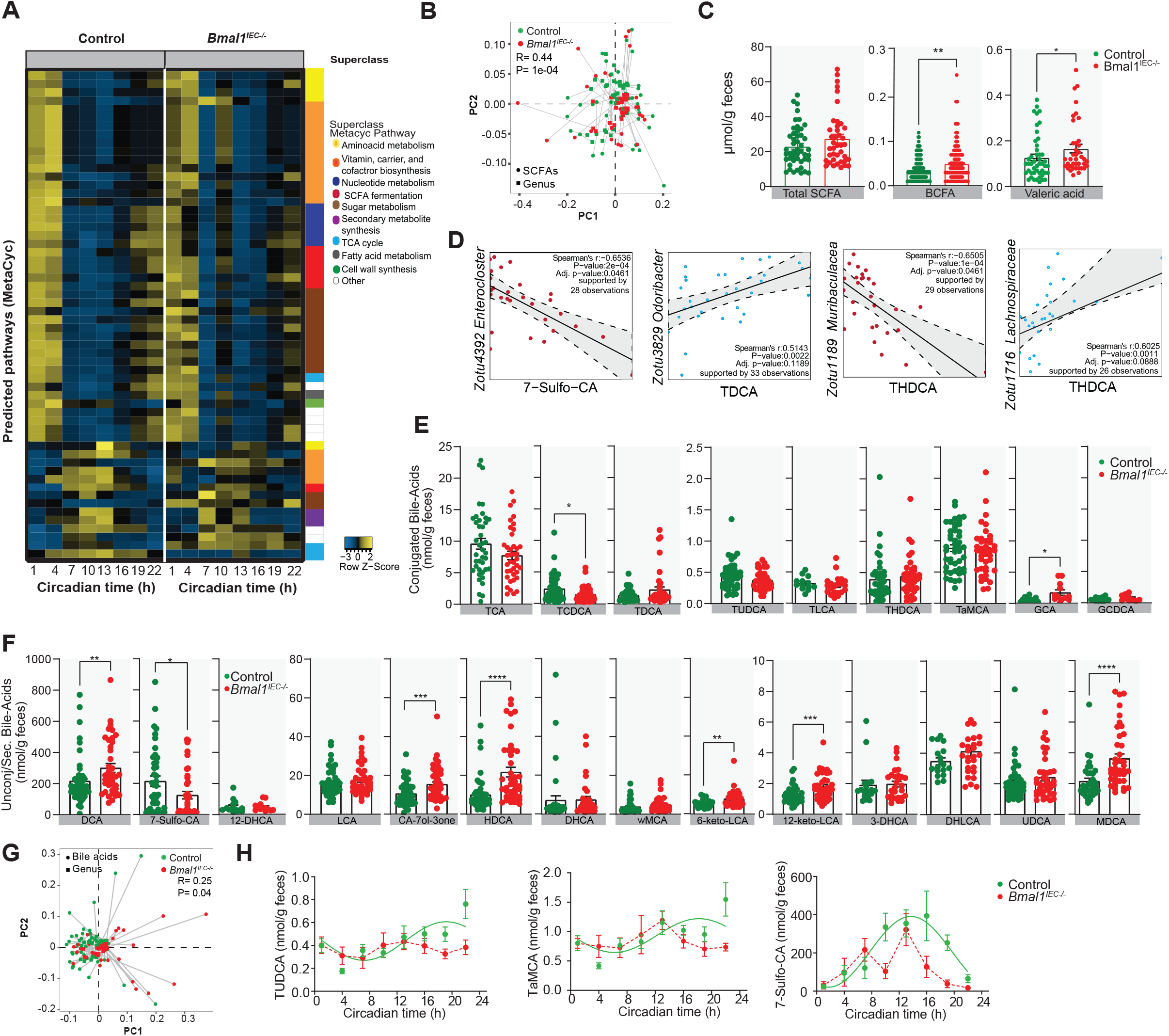
Metabolic functioning of intestinal clock-controlled bacteria. (A) Heatmap of MetaCyc Pathways predicted using PICRUST2.0 on intestine clock-controlled zOTUs rhythmic in control (left) and arrhythmic in *Bmal1^IEC-/-^* (right) mice. Pathways are colored according to their sub-class. (B) Procrustes analyses (PA) of fecal microbiota and SCFA levels. The length of the line is proportional to the divergence between the data from the same mouse. (C) SCFA concentrations in feces. (D) and Spearman correlation (p-value ≤ 0.05 and R ≤ −0.5; red or R ≥ 0.5; blue) between SCFA and gut controlled bacteria taxa. (E) Conjugated and secondary (F) fecal BAs levels. (G) PA as described in (D) with fecal bile-acid (BA) levels. (H) Circadian profiles of BAs. Significant rhythms are illustrated with fitted cosine-regression (solid line); data points connected by dotted lines indicate no significant cosine fit curves (p-value > 0.05) and thus no rhythmicity. n = 5-6 mice/time point/genotype. Control (green) and *Bmal1^IEC-/-^* (red). Data are represented as mean ± SEM. Significance * p ≤ 0.05, ** p ≤ 0.01, *** p ≤ 0.001, **** p ≤ 0.0001 (Mann-Whitney U test). BCFA=branched-chain fatty acids. Cholic acid (CA), a-Muricholic acid (aMCA), b-Muricholic acid (bMCA), Taurocholic acid (TCA), Taurochenodeoxycholic acid (TCDCA), Tauroursodeoxycholic acid T(aTuUroDhCyAo)d, eoxycholic acid (THDCA), Taurolithocholic acid (TLCA), Taurodeoxycholic acid (TDCA), Tauro-a-Muricholic acid (TaMCA), Glycochenodeoxycholic a c(GidCDCA), Glycocholic acid (GCA), Deoxycholic acid (DCA), Lithocholic acid (LCA), y-Muricholic acid (y-MCA), 12-Dehydrocholic acid (12-DHCA), 12-Ketolithocholi cacid (12-ke-to-LCA), 3-Dehydrocholic acid (3-DHCA), 6-Ketolithocholic acid (6-keto-LCA), 7-Dehydrocholic acid (7-DHCA), 7-Sulfocholic acid (7-sulfo-CA), Allocho laiccid (ACA), Cholic acid-7ol-3one (CA-7ol-3one), Ursocholic acid (UCA), Dehydrolithocholic acid (DHLCA), Hyodeoxycholic acid (HDCA), Murideoxycholi c(MaDciCdA), Ursodeoxycholic acid (UDCA).

### Transfer of arrhythmic intestinal clock-controlled microbiota disturbs GI homeostasis

To further test the physiological relevance of intestinal clock-driven rhythmic microbial composition and its functionality on host physiology, GF C57BL/6 mice were colonized with cecal content from *Bmal1^IEC-/-^* or control mice (n=4) (**Fig. 6A**). Importantly, ~76 % of bacteria from the control and *Bmal1^IEC-/-^* donor, were transferred into the recipients (**Suppl. Fig. 3E**). Reduced richness was observed in mice after microbiota transfer (**Suppl. Fig. 3F**), in line with previous reports ^38^. Interestingly, microbiota composition of recipient wild-type hosts significantly differed depending on the genotype of the donor (**Fig. 6B; Suppl. Fig. 3G,H**). Most of the transferred taxa belonged to Firmicutes, whose rhythmicity strongly depends on a functional intestinal clock, whereas abundances in Bacteroidetes were highly suppressed (**Fig. 6C; Suppl. Fig. 3G**). Importantly, lack of rhythmicity was transferred to mice receiving microbiota from *Bmal1^IEC-/-^* mice (**Fig. 6D, E**). In particular rhythmicity of zOTUs belonging to e.g. *Alistipes*, shown to be driven by the gut clock (**Fig. 3F**) and maintained circadian rhythmicity in the host after transfer from control donors, lack rhythmicity in mice receiving microbiota from *Bmal1^IEC-/-^* mice (**Fig. 6E**). Interestingly, after transfer of *Bmal1^IEC-/-^* - associated arrhythmic microbiota several bacterial-derived metabolites, including BCFA as well as the secondary bile ketolitocholic acids were significantly upregulated and the primary BA β-muricholic acid (b-MCA) was significantly downregulated **(Fig. 6F)**, similar to observations in made in donor mice **(Fig 5C,F, Suppl. Fig. 3A-D).** Moreover, altered expression of intestinal clock genes in jejunum and colon was observed after transfer of arrhythmic microbiota in comparison to recipient mice receiving rhythmic microbiota (**Suppl**. **Fig. 4A**). In addition the expression of genes involved in host-microbe interaction, including *Arg2* and *Tlr4*, significantly differed depending on the genotype of the donor (**Fig. 6G**). Arrhythmic microbiota transfer induced the expression of *Il33 and NfkB* both known to be involved in intestinal inflammatory responses ^39,40^ and reduced the expression of *Ang4* and *Hdac3* (**Fig. 6G**), which are known to integrate microbial and circadian cues relevant for inflammatory and metabolic intestinal functions ^33,41,42,31,35^. Although mesenteric lymph nodes and colon weights of mice associated with arrhythmic microbiota were undistinguishable from controls, and histology scores were unaffected, we noticed an increase in spleen and jejunum weight (**Fig. 6H, Suppl. Fig. 4B, C**), indicating a role of microbial rhythmicity on host intestinal homeostasis. Indeed, transfer of arrhythmic microbiota altered immune cell recruitment to the lamina propria. For example, an increase in dendritic cells (CD11c+) was detected in the small intestine whereas recruitment of T cells (CD3+CD4+, CD3+CD8+) and dendritic cells (CD11c+) to the lamina propria of the colon was decreased (**Fig. 6I, Suppl. Fig. 4D**). Similar trends in immune cell recruitment and intestinal gene expression were observed in *Bmal1^IEC-/-^* mice in SPF conditions **(Suppl. Fig. 4E, F)**, suggesting that intestinal-clock controlled microbes influence gut immune homeostasis. Importantly, alterations found following transfer of *Bmal1^IEC-/-^* -associated arrhythmic microbiota were not observed after transfer of microbiota from control mice kept in either LD, DD or starvation conditions, highlighting the physiological relevance of gut-clock-controlled microbiota **(Suppl. Fig. 4G-I).**

**Figure 6.**
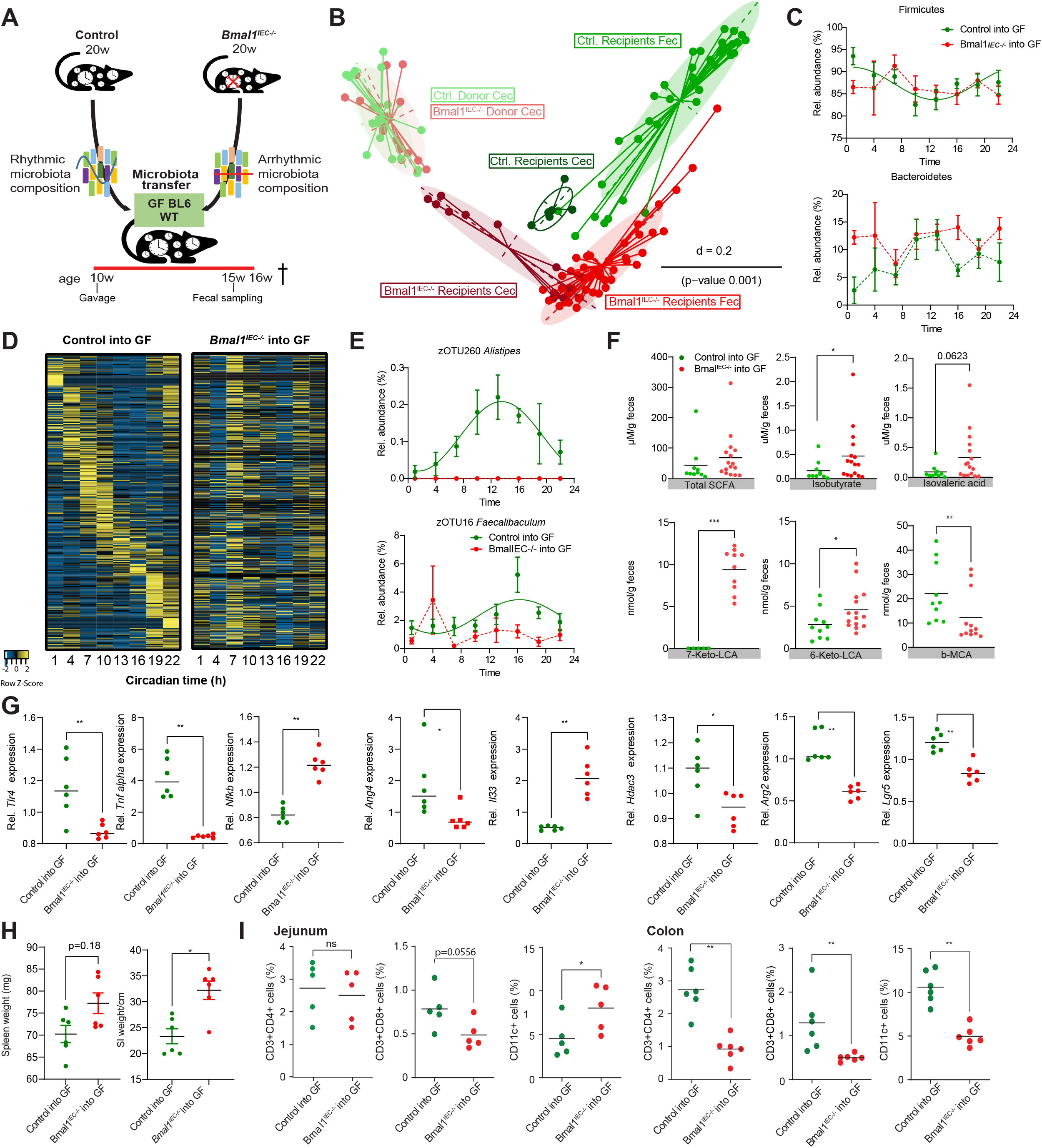
Intestinal clock-controlled rhythmic microbiota is essential for intestinal homeostasis. (A) Schematic illustration of transfer experiments with mixture of cecal microbiota obtained from *Brnal1^IECfl/fl^* control and *Bmal1^IEC-/-^* donors (n=4) into 10 weeks old germ-free BL6 wild-type recipient mice till sacrifice at 16 weeks (16w). Fecal profiles were collected 5 weeks after gavage at the age of 15 weeks (15w). (B) PCoA plot of GUniFrac distances of donor (CT13) and recipient mice (n=6/genotype/time point) after 5 weeks of transfer. (C) Diurnal profiles of fecal microbiota at phylum level. (D) Heatmap depicting the relative abundance of zOTUs ordered by their cosine-regression peak phase according to the recipient controls. On the left the first two columns indicate donor zOTU abundance. (E) Diurnal profile of relative abundance of example zOTUs. (F) Fecal SCFA and BAs concentrations. (G) Relative gene expression of Tlr4, Arg2, Ang4, Tnfa, Il33, Nfkb, Hdac3, Lgr5 in the proximal colon of recipient mice. (H) Organ weights of recipient mice after receiving control (green) or *Bmal1^IEC-/-^* (red) cecal microbiota. (H) Frequency of CD3+CD4+, CD3+CD8+, CD11c+ cells in jejunum rhythms are illustrated with fitted cosine-regression; data points connected by dotted lines indicate no significant cosine fit curves (p-value > 0.05) and thus no rhythmicity. n = 6 mice/ genotype. Data are represented as mean ± SEM. Significance * p ≤ 0.05, ** p ≤ 0.01 (Mann-Whitney U test).

Together, these results indicate that deletion of intestinal clock function caused loss of rhythmicity in the majority of bacterial taxa, alters microbial functionality and resets the host’s intestinal homeostasis.

## Discussion

Diurnal microbial oscillations in the major phyla and families have previously been demonstrated in humans and mice by us and others ^9–13^. Here we provide evidence that rhythmicity of the majority of these taxa persists even in the absence of external timing cues, demonstrating the endogenous origin of fecal bacterial circadian rhythms. In contrast, strongly attenuated microbial rhythms in cecal and ileal content were found in mice kept in DD ^43^. Differences in the amount and type of oscillating microbes between studies may be explained by variations in microbiota composition of the mouse lines, microbial ecosystem between facilities and niches within the GI tract. Moreover, our comparison analysis indicates that the discrepancy in total amounts of oscillating taxa between our study and previous studies^9,10^ is likely due to the different methods used to classify groups of closely related taxa. The comparability between studies may additionally be impacted by methodological differences, such as differences in the 16S rRNA variable target gene region used for amplicon sequencing (here: V3-V4, ^43^: V4, ^10^: V1-V2, ^21^.: V1-V3, ^44^: Metatranscriptome) as well as the sampling intervals (here: 3h/day, ^43^: 6h/day, ^10^: 4h/day).

Our results obtained from constant environmental conditions suggest that circadian microbiota regulation is primarily rooted in host or bacterial intrinsic circadian mechanisms. Indeed, using IEC-specific *Bmal1*-deficient mice, we provide the first demonstration for a dominant role of intestinal clocks in driving circadian microbiota composition. Lack of *Bmal1* in IECs led to dramatic loss of microbiota rhythmicity, predominantly belonging to the phylum Firmicutes, which was independent of whether relative or quantitative analyses was used. Of note, both microbiota analyses in this study did not always yield identical results, highlighting the importance of both analyses to interpret circadian microbiota composition. Microbiota composition is commonly analyzed by relative abundance; yet this analysis may exaggerate microbial rhythmicity due to masking by highly abundant taxa ^13^. Arrhythmicity in taxa (e.g. *Lactobacillaceae, Odoribacteraceae* and *Lachnospiraceae*) was previously documented in mice lacking *Bmal1* or *Per1/2* tissues-wide ^10,13^. Loss of oscillations of the same taxa among others was observed in *Bmal1^IEC-/-^* mice, identifying the IEC-clock to generate rhythmicity of these bacteria. This prominent role of the intestinal clock in driving microbial rhythmicity was further validate on data obtained in LD condition. Few of the remaining bacteria which were unaffected by IEC clock-deficiency, lost circadian rhythmicity upon starvation. Some of these taxa have previously been reported to be influenced by food availability ^11^. A prominent role for the timing of food intake on microbial rhythmicity was frequently suggested ^9–11^. However, we demonstrate that even in the absence of food, rhythmicity of dominant bacteria persisted, thus rhythmic food intake is not required to drive microbiota oscillations. Accordingly, in mice supplied with constant intravenous parenteral nutrition cecal microbiota composition oscillated ^9^. Importantly, the majority of microbial rhythms was abolished in *Bmal1^IEC-/-^* mice even though these mice show rhythmic food intake behavior. These results further highlight that IEC clocks are the dominant driver whereas environmental changes as well as food-intake are mere modulators of microbial rhythms.

Mechanisms how intestinal clocks regulate microbiota rhythms likely involve local epithelial-microbe interactions and immune functions, including pattern recognition receptors, such as Toll-Like Receptors (TLRs), Aryl hydrocarbon receptors and nuclear receptors as well as antimicrobial peptide production and mucus secretion, all previously found to oscillate diurnally ^32,33,46^. Accordingly, we found IEC clocks to be essential for rhythmic expression of related genes, such as *Tlr2, Ang4, Muc2, Nfil3* and *Hdac3*, which are engaged in bidirectional microbe-host communication. For example, IEC-specific deletion of *Hdac3* or loss of *Tlr2* altered microbial composition and influenced intestinal homeostasis ^47,48^. Furthermore, *Muc2* can affect microbiota composition by allowing bacteria to metabolize mucin glycans ^49^, which through cross-feeding affects the abundance of other microbial taxa ^50^. However, future studies are required to functionally investigate intestinal clock mechanism controlling microbiota composition.

Circadian disruption due to life style has been correlated to microbiota dysbiosis and human health (reviewed by ^51^). Recently, we found a functional link between arrhythmicity of microbiota and the development of obesity and T2D ^12^. Here we demonstrate intestinal-clock dependent rhythmicity of taxa involved in SCFA fermentation, such as *Lachnospiraceae, Ruminococcaceae* and *Odoribacteraceae*, lactate acid producing bacteria e.g. *Lactobacillaceae*, mucus foragers, such as *Muribaculaceae, Rickenellaceae* and *Lachnospiraceae* ^36,52,53^. Moreover, taxa capable to convert bile acids (BAs), including *Lactobacillus* and *Ruminococcus*^54^ are regulated by intestinal circadian clocks. Consequently, IEC clocks likely influence microbiota function. Indeed, functional analysis based on 16S rRNA gene data revealed pathways such as SCFA fermentation, amino-acid and carbohydrate metabolism are associated to intestinal clock-controlled taxa. Many among the identified pathways were shown to diurnally oscillate in healthy individuals, to lose rhythmicity in clock deficient mice, and are associated with host’s health ^10,12,55^. Targeted metabolite analysis further confirmed that arrhythmicity in intestinal clock-controlled taxa is reflected in alterations of key metabolites involved in lipid signaling pathways such as SCFAs and BAs. Both of these microbiota-related metabolites are known to diurnally oscillate and impact, among others host immune and metabolic functioning ^9,56–60^. Notably, high levels of secondary BAs, e.g. DCAs and HDCAs identified in *Bmal1^IEC-/-^* mice, were also found in subjects with metabolic disorders including obesity and T2D ^61–64^. Moreover, TUDCA lost circadian rhythmicity in *Bmal1^IEC-/-^* mice and was found to ameliorate insulin sensitivity in both obese mice and humans ^65,66^. In addition, accumulation of BAs within the host has been associated with several GI diseases. For example, increased levels of the sec. BA DCA and taurine-conjugated BAs were linked to colorectal cancer in mice (reviewed by Jia and colleagues ^67^). These results support the hypothesis that a functional intestinal clock balances GI health by driving microbial function.

Transfer experiments provide direct evidence for the physiological relevance of intestinal clock-controlled microbiota and their functions. Microbial rhythmicity and arrhythmicity depending on the donor genotype induced molecular and physiological changes in the healthy hosts. Notably, a functional intestinal clock in recipients seems to require longer than 5 weeks to restore microbial rhythmicity, suggesting strong microbe-host clock interactions. Indeed, this was supported by altered clock gene expression in recipients receiving arrhythmic microbiota and is in accordance with a report indicating a role of microbial rhythmicity in programing host transcriptome rhythmicity ^24^. Additionally, the alteration of microbial metabolite production in recipients, which reflect the results obtained from donor mice, e.g. upregulated BCFAs and sec. BAs. may have influenced intestinal clock functions, since a direct impact on rhythmicity in intestinal epithelial cells and subsequent metabolic responses in the host has been provided previously ^9,49^.

Transfer of arrhythmic microbiota from intestinal clock-deficient mice altered several genes involved in inflammation, antimicrobial peptide production and intestinal epithelial functioning including *Tnf-α, Tlr2, Lgr5, Ffar2, Hdac3* and *Nf-kB* ^39–41,68–71^. Changes in the expression of these genes due to reception of arrhythmic microbiota might have caused the observed immune phenotype in recipients. For example, *Tlr2* senses the presence of microbe-associated molecular patterns and is capable to regulate the host’s immune system against pathogenic infiltration by regulating *Tnf-α* cytokine production of CD8+ T-cells ^72,73^. Moreover, *Hdac3* and *Il33* also plays an important role in microbe-host crosstalk ^47,74^. Mouse models lacking HDAC3 show high susceptibility to DSS-induced inflammation, partly by activation of NF-kb, the latter also observed in our mouse model ^47,75^. Consequently, altered regulation of *Tlr2, Tnf-α, Hdac3*, and *IL-33* expression observed in mice receiving arrhythmic microbiota, likely disturbed gastrointestinal homeostasis. Indeed, enhanced lymphoid tissues weight and suppressed recruitment of T-cell and dendritic cell populations to the lamina propria within the colon was found in GF mice receiving arrhythmic microbiota from intestinal clock-deficient mice. Immune cell recruitment represents an important aspect in the gastrointestinal immune defense (reviewed by ^76^). Interestingly, we observed similar alterations and microbial-derived metabolite changes in recipients and SPF donors, indicating that microbiota might transfer the phenotype from the donor to recipients. Discrepancies between the immune phenotype observed in donor and recipient mice might be caused by genetic loss of Bmal1 in the intestine of donors and feedback from arrhythmic microbiota on intestinal tissue of rhythmic recipients. In contrast, transfer of microbiota from control mice kept in LD, DD or in starving conditions did not cause the same alterations in immune cell recruitment, gene expression or microbial metabolites levels. These results provide the first evidence that intestinal clock-controlled rhythmic gut bacteria are crucial for a balanced intestinal immune homeostasis, and likely influence the immune response to pathogens, infection and inflammation.

Taken together, a functional intestinal clock represents a key element to maintain gastrointestinal health by driving rhythmicity of gut bacteria and microbial products. Since intestinal clock functions effect bacterial taxa required for a balanced immune defense, it remains to be studied whether arrhythmicity of intestinal clock-controlled taxa is causal for the development of gastrointestinal diseases. In the scope of previous associations of arrhythmic microbiota and their products with T2D and IBD ^12,77^, our data highlight the relevance to further investigate intestinal clock mechanisms driving bacterial rhythmicity for human health.

## ADDITIONAL INFORMATION (CONTAINING SUPPLEMENTARY INFORMATION LINE (IF ANY) AND CORRESPONDING AUTHOR LINE)

## ACKNOWLEDGEMENT

The Technical University of Munich provided funding for the ZIEL Institute for Food & Health, animal facility support, technical assistance and support for 16S rRNA gene amplicon sequencing. Johanna Bruder provided assistance with animal experiments and preliminary data collection.

## AUTHOR CONTRIBUTIONS

SK conceived and coordinated the project. MH, BA performed mouse experiments and fecal samples collection. BA, MH and YN provided tissue samples and performed gene expression analysis. SK and MH analyzed activity and food intake behavior. MH, BA, SR and SK performed 16S rRNA gene sequencing and bioinformatics analysis. BA and MH analyzed predicted microbial functionality. YN performed FACS analyses. MH and KK performed targeted metabolomics and data analysis. SK supervised the work and data analysis. SK and DH secured funding. MH, BA, DH and SK wrote the manuscript. All authors reviewed and revised the manuscript. MH and BA contributed equally to this work.

## MATERIALS & CORRESPONDENCE

Correspondence to Dr. Silke Kiessling, Chair of Nutrition and Immunology, Technical University of Munich, Gregor-Mendel-Str. 2, 85354 Freising, Germany.

## METHODS

### Ethics Statement

Experiments were conducted at Technical University of Munich in accordance with Bavarian Animal Care and Use Committee (TVA ROB-55.2Vet-2532.Vet_02-18-14).

### Mouse models

#### Bmal1^IEC tg/wt^ and Bmal1^IECfl/fl^ mouse generation

Male epithelial intestinal epithelial cell-specific *Bmal1* knock-out (*Bmal1fl/fl* x Villin CRE/wt; referred to as *Bmal1^IEC-/-^*) mice and their control littermates (*Bmal1fl/fl* x Villin wt/wt; referred to as *Bmal1^fl/fl^*) on a genetic C57BL/6J background were generated as previously described ^78^. Breeding was performed by crossing *Bmal1fl/fl* x Villin CRE/wt with *Bmal1fl/fl* x Villin wt/wt. Unless otherwise stated, mice were kept in LD 12:12 cycles (300 lux), with lights turned on at 5am (*Zeitgeber* time (ZT0) to 5pm (ZT12)). All mice were single housed at the age of 8 weeks in running wheel-equipped cages with ad libitum access to chow and water and under specific-pathogen free (SPF) conditions according the FELASA recommendation. In order to minimize cage-related bias in microbiota composition ^79^, littermates and litters of comparable age from as few as possible breeding pairs and cages were selected.

#### Behavior analysis

All male mice, unless stated otherwise, were individually housed in cages with running wheels. Handling and activity measurements during experiments were performed as described ^80^. Wheel-running activity was analyzed using ClockLab software (Actimetrics). The last 10-14 days of each condition were used to determine the period (tau, calculated using a X^2^ periodogram and confirmed by fitting a line to the onsets of activity), the duration of the active period (alpha), the amount of activity and the subjective day/night activity ratio (where the subjective day under DD conditions is the inactive period between the offset of activity and the onset of activity and the subjective night is the active period between the onset of activity and the offset of activity).

#### Light-dark (LD) and constant darkness (DD) conditions and fecal sample collection

Male *Bmal1^IEC-/-^* and their control littermates *Bmal1^IECfl/fl^* were maintained under LD cycle for 2 weeks (age 8-10 weeks), switched to a DD cycle for 2 more weeks (age 10-12 weeks), kept in constant light (LL) for an additional 2 weeks (age 12-14w) and finally returned back to LD till the age of 18-20 weeks before sacrifice. Average daily food intake was measured over 5 consecutive days in the second week of the above indicated light conditions. Of note, Fecal sample collection in darkness was performed after adjusting for each mouse’s individual free-running period (**Suppl. Fig. 2A**). For example, the activity period varied by ~0.5 hours per day within mice of the same genotype and consequently accumulates to a ~7 hour phase difference between individual mice after 2 weeks in DD. Fecal collection in LD was performed according to normal *Zeitgeber* time. Samples were collected every 3 hours over the course of a 24h-day (**Fig. 1A, Suppl. Fig. 2A**).

#### Food deprivation

11-week-old *Bmal1^IEC-/-^* and control male mice with *ad libitum* food intake were used for fecal sample collection. At the age of 12-13 weeks control littermates from *Bmal1^IEC-/-^* mice were starved beginning at CT1 on the 1^st^ day in DD. Fecal samples were collected as detailed above starting at CT13 on the 2^nd^ day in DD (after 13 hours of starvation) till CT10 (after 34 hours of starvation) at the indicated time points (**Fig. 2A**).

#### Food intake pattern and analysis

The diurnal food intake pattern of individually housed mice was recorded using an automated monitoring system (TSE LabMaster Home Cage Activity, Bad Homburg, Germany) at the age of 10-12 weeks. Mice were habilitated for 3 days. Then data were collected for a full 24-hours profile at the 4^th^ day. Cumulative food intake was recorded through high precision balances connected to the food baskets. Food basket weight were summed up in 1 min intervals. Total food intake was calculated as 1^st^ derivative of cumulative food intake and consumption was summed up at intervals of 1 hour. One-minute intervals where the weight loss was directly followed by a similar weight gain in food baskets within a period of 3 minutes were excluded. Circadian food intake of individually housed mice was recorded in the second day of darkness by weighing the food every 3 hours.

#### Complete gastrointestinal transit time

Complete GI transit time (GITT) was measured by administering natural carmine red (6%, SigmaAldrich) dissolved in 0.5% methylcellulose (Sigma-Aldrich) by gavage (100ul) according to ^81^. 16-18 week-old male *Bmal1^IEC-/-^* and *Bmal1^fl/fl^* mice were starved 6h prior to gavage. Time of gavage was considered T0. In case of presence, fecal pellets were taken out of the cage every 10 minutes and checked for red color. GITT was registered as the time between T0 and the time of presence of carmine red in the fecal pellets.

#### Tissue collection in SPF conditions

All animals were sacrificed by cervical dislocation at the age of 18-20 weeks in the second day of darkness at the indicated circadian times (CT) or in LD12:12 conditions, in 4 hour intervals starting at 1 hour after lights off (ZT1). In constant darkness samples were collected at the indicated time points used for control mice in LD. Eyes were removed prior to tissue dissection. Tissues were harvested and snap frozen on dry ice and stored in −80 degrees.

#### Gene expression analysis (qRT-PCR) Quantitative real-time PCR

RNA was extracted from snap frozen tissue samples with Trizol reagent. cDNA was synthesized from 1000ng RNA using cDNA synthesis kit Multiscribe RT (Thermofischer Scientific). qPCR was performed in a Light Cylcer 480 system (Roche Diagnostiscs, Mannheim, Germany) using Universal Probe Library system according to manufacturer’s instructions. For genes expression the following primers and probes were used: Brain and Muscle ARNT-Like 1 (*Bmal1*) F 5’-ATTCCAGGGGGAACCAGA-’ R 5’-GGCGATGACCCTCTTATCC-3’ Probe 15, Period 2 (*Per2*) F 5’-TCCGAGTATATCGTGAAGAACG-3’ R 5’-CAGGATCTTCCCAGAAACCA-3’ probe 5, Nuclear receptor subfamily 1 group D member 1 (*Reverba*) F 5’-AGGAGCTGGGCCTATTCAC-3’ R 5’-CGGTTCTTCAGCACCAGAG-3’ probe 1, Toll-like receptor 2 (*Tlr2*) F-5’-GGGGCTTCACTTCTCTGCTT-3’ R 5’-AGCA TCCTCTGAGATTTGACG-3’ probe 50, Angiogenin 4 (*Ang4*) F 5’-CCCCAGTTGGAGGAAAGC-3’ R 5’-CGTAGGAATTTTTCGTACCTTTCA-3’ probe 106, Mucin 2 (*Muc2*) F 5’-GGCAGTACAAG AACCGGAGt-3’ R 5’-GGTCTGGCAGTCCTCGAA-3’ probe 66, Histone Deacetylase 3 (*Hdac3*) F 5’-GAGAGGTC CCGAGGAGAAC-3’ R 5’-CGCCATCATAGAACTCATTGG-3’ probe 40, Tumor necrosis factor alpha (*Tnfa*) F 5’-TGCCTATGTCTCAGCCTCTTC-3’ R 5’-GAGGCCATTTGGGAACTTCT-3’ probe 49, Leucine Rich Repeat Containing G Protein-Coupled Receptor 5 (*Lgr5*) F 5’-CTTCACTCGGTGCAGTGCT-3’ R 5’-CAGCCAGCTACCAAATAGGTG-3’ probe 60, Toll like receptor 4 (*Tlr4*) F 5’-GGACTCTGATCATGGCACTG-3’ R 5’-CTGATCCATGCATTGGTAGGT-3’ probe 2, Nuclear factor kappa-light-chain-enhancer of activated B cells (*Nfkb*) F 5’-CCCAGACCGCAGTATCCAT-3’ R 5’-GCTCCAGGTCTCGCTTCTT-3’ probe 47. RNA abundance was normalized to the housekeeping gene Elongation factor 1-alpha (*Ef1a*) F 5’-GCCAAT TTCTGGTTGGAATG-3’ R 5’-GGTGACTTTCCATCCCTTGA-3’ probe 67. For Interleukin33 (*Il33*) gene was used syber green to run the gene using the following primer F 5’-GAACATGAGTCCCATCAAAG-3’ R 5’-CAGCTGGTTATCTTTTACTCC-3’ and RNA abundance was normalized to the housekeeping gene *Ef1a* F 5’-GCCAAT TTCTGGTTGGAATG-3’ R 5’-GGTGACTTTCCATCCCTTGA-3’.

#### High-Throughput 16S Ribosomal RNA (rRNA) Gene Amplicon Sequencing Analysis

Genomic DNA was isolated from snap-frozen fecal pellets according to a modified protocol of Godon and colleagues ^82^, as previously described ^12^. DNA NucleoSpin gDNA columns (Machery-Nagel, No. 740230.250) were used for DNA purification. In a two-step PCR the V3-V4 region (using the primers 341F-ovh and 785r-ov) of the 16S rRNA gene was amplified from 24 ng DNA. After pooling, the multiplexed samples were sequenced on an Illumina HiSeq in paired-end mode (2×250 bp) using the Rapid v2 chemistry, in accordance with ^12^. Two negative controls, consisting of DNA stabilizer without stool, were used for every 45 samples to control for artifacts and insure reproducibility. High-Quality sequences of read counts > 5000 were used for 16s rRNA data analysis. Reads FASTQ files were consequently processed using an in-house developed NGSToolkit (Version Toolkit 3.5.2_64) based on USEARCH 11 ^83^. A trim score of 5 was used on the 5’ end and 3’end for the R1 and R2 read followed by chimera removal ^84^ using the FASTQ mergepair script of USEARCH ^83^. Quality filtered reads were merged, deduplicated, clustered and a denoised clustering approach was applied to generate zero-radius operational taxonomic units (zOTUs) ^83,85^. Using zOTUs analysis provide the best possible resolution considering the 16s rRNA sequencing, by correcting the sequencing error (denoising step) and then identifying each unique sequence (100% similarity) as a different microbial strain. On the other hand, previously OTU analysis was used, which is based on merging different strains which has more than 97% similarity into one single OTU, and assign it to one microbial strain^45^. Taxonomic assignment was performed with the EZBiocloud database^86^. Notably, For Suppl. Fig 1L Taxa were assigned to Greengenes database in order to be able to compare our results with the data obtained in the study by Thaiss and colleagues ^10^. Data was further analyzed with the R-based pipeline RHEA ^87^. Phylogenetic trees are generated by a maximum likelihood approach, which was performed on an alignment generated by MUSCLE with the software MegaX ^88^. Trees were visualized and annotated with the use of the online tool EvolView (http://www.evolgenius.info/evolview/)^89^. Spike-in of 12 artificial DNA standards mimicking 16S rRNA genes (a surrogate for bacterial numbers) were used to determine the quantitative copy numbers of rRNA genes per gram of fecal sample between samples. Briefly, the same amount of artificial DNA (6ng) was added to each weighted fecal sample before DNA extraction. After sequencing as described above, FASTQ files were mapped against the spike FASTA sequences (using bowtie2), removing the spike reads and generating a new FASTQ file. By comparing the spike sequencing reads to the fecal bacterial reads; we calculate the quantitative number of 16S rRNA gene copies per gram of sample. The copy number of 16S rRNA gene is proportional to the number of bacteria present in a sample. Thus, this approach enables estimation of microbial abundances relative between samples, suitable for comparative analysis according to Tourlousse et al. 2017 ^23^.

### Targeted metabolite analyses

#### Sample preparation for targeted metabolite analyses

Approximately 20 mg of mouse fecal pellet was weighed in a 2 mL bead beater tube (CKMix 2 mL, Bertin Technologies, Montigny-le-Bretonneux, France) filled with 2.8 mm ceramic beads. 1 mL of methanol-based dehydrocholic acid extraction solvent (c=1.3 μmol/L) was added as an internal standard for work-up losses. Fecal samples were extracted with a bead beater (Precellys Evolution, Bertin Technolgies) supplied with a Cryolys cooling module 3 times each for 20 seconds with 15 seconds breaks in between at 10.000 rpm.

#### Targeted bile acid measurement

20 μL of isotopically labelled bile acids (ca. 7 μM each) were added to 100 μL of sample extract. Targeted bile acid measurement was performed using a QTRAP 5500 triple quadrupole mass spectrometer (Sciex, Darmstadt, Germany) coupled to an ExionLC AD (Sciex, Darmstadt, Germany) ultrahigh performance liquid chromatography system. A multiple reaction monitoring (MRM) method was used for the detection and quantification of the bile acids. An electrospray ion voltage of −4500 V and the following ion source parameters were used: curtain gas (35 psi), temperature (450 °C), gas 1 (55 psi), gas 2 (65 psi), and entrance potential (−10 V). The MS parameters and LC conditions were optimized using commercially available standards of endogenous bile acids and deuterated bile acids, for the simultaneous quantification of selected 28 analytes. For separation of the analytes a 100 × 2.1 mm, 100 Å, 1.7 μm, Kinetex C18 column (Phenomenex, Aschaffenburg, Germany) was used. Chromatographic separation was performed with a constant flow rate of 0.4 mL/min using a mobile phase consisted of water (eluent A) and acetonitrile/water (95/5, v/v, eluent B), both containing 5 mM ammonium acetate and 0.1% formic acid. The gradient elution started with 25% B for 2 min, increased at 3.5 min to 27% B, in 2 min to 35% B, which was hold until 10 min, increased in 1 min to 43% B, held for 1 min, increased in 2 min to 58% B; held 3 min isocratically at 58% B, then the concentration was increased to 65% at 17.5 min, with another increase to 80% B at 18 min, following an increase at 19 min to 100% B which was hold for 1 min, at 20.5 min the column was equilibrated for 4.5 min at starting. The injection volume for all samples was 1 μL, the column oven temperature was set to 40 °C, and the auto-sampler was kept at 15 °C. Data acquisition and instrumental control were performed with Analyst 1.7 software (Sciex, Darmstadt, Germany) as previously described ^90^. Briefly, a standard curve (0.5 nM to 15000 nM, 16 points), linear regression and weighting type of 1/x was used to calculated BA concentrations ^90^. BAs measured are Cholic acid (CA), chenodeoxycholic acid (CDCA), a-Muricholic acid (aMCA), b-Muricholic acid (bMCA), Taurocholic acid (TCA), Taurochenodeoxycholic acid (TCDCA), Tauroursodeoxycholic acid (TUDCA), Taurohyodeoxycholic acid (THDCA), Taurolithocholic acid (TLCA), Taurodeoxycholic acid (TDCA), Tauro-a-Muricholic acid (TaMCA), Glycochenodeoxycholic acid (GCDCA), Glycocholic acid (GCA), Deoxycholic acid (DCA), Lithocholic acid (LCA), y-Muricholic acid (y-MCA), 12-Dehydrocholic acid (12-DHCA), 12-Ketolithocholic acid (12-keto-LCA), 3-Dehydrocholic acid (3-DHCA), 6-Ketolithocholic acid (6-keto-LCA), 7-Dehydrocholic acid (7-DHCA), 7-Sulfocholic acid (7-sulfo-CA), Allocholic acid (ACA), Cholic acid-7ol-3one (CA-7ol-3one), Ursocholic acid (UCA), Dehydrolithocholic acid (DHLCA), Hyodeoxycholic acid (HDCA), Murideoxycholic acid (MDCA), Ursodeoxycholic acid (UDCA).

#### Targeted short-chain fatty acid measurement

For the quantitation of short-chain fatty acids (SCFAs) the 3-NPH method was used ^91^. Briefly, 40 μL of the fecal extract and 15 μL of isotopically labeled standards (ca 50 μM) were mixed with 20 μL 120 mM EDC HCl-6% pyridine-solution and 20 μL of 200 mM 3-NPH HCL solution. After 30 min at 40°C and shaking at 1000 rpm using an Eppendorf Thermomix (Eppendorf, Hamburg, Germany), 900 μL acetonitrile/water (50/50, v/v) was added. After centrifugation at 13000 U/min for 2 min the clear supernatant was used for analysis. The same system as described above was used. The electrospray voltage was set to −4500 V, curtain gas to 35 psi, ion source gas 1 to 55, ion source gas 2 to 65 and the temperature to 500°C. The MRM-parameters were optimized using commercially available standards for the SCFAs. The chromatographic separation was performed on a 100 × 2.1 mm, 100 Å, 1.7 μm, Kinetex C18 column (Phenomenex, Aschaffenburg, Germany) column with 0.1% formic acid (eluent A) and 0.1% formic acid in acetonitrile (eluent B) as elution solvents. An injection volume of 1 μL and a flow rate of 0.4 mL/min was used. The gradient elution started at 23% B which was held for 3 min, afterward the concentration was increased to 30% B at 4 min, with another increase to 40%B at 6.5 min, at 7 min 100% B was used which was hold for 1 min, at 8.5 min the column was equilibrated at starting conditions. The column oven was set to 40°C and the autosampler to 15°C. Data acquisition and instrumental control were performed with Analyst 1.7 software (Sciex, Darmstadt, Germany). Concentrations of SCFA were calculated with the use of linear regression using a standard curve (ranging 0.0004 mM to 2 mM (12 points), SCFAs measured are Acetate, Propionate, Butyrate, Valeric acid, Desaminotyrosine and the Branched-chain Fatty acids (Isobutyric acid, 2-Methylbutyric acid and Isovaleric acid).

#### Targeted metabolite analyses

MultiQuant 3.0.3 Software (AB Sciex) was used to integrate the data and calculate the concentration. Isotopically labelled standards were used for SCFA quantitation. For bile acid quantitation, we used Dehydrocholic acid as internal standard to correct for losses during sample preparation and isotopically labelled references to correct for ionization effects during measurement, according to the paper of Sinah et al. 2021 ^90^. For comparison between two groups, Mann-Whitney U test was used to test for statistical significance. Metabolite-microbiota correlation analyses was performed on relative abundance zOTU level within the rhythmic in male *Bmal1^fl/fl^*, but not *Bmal1^IEC-/-^* samples with at least a 30% prevalence. Spearman correlation and adjusted p-values between targeted metabolomics and zOTUs were calculated using the *rcor(*) function in R. Correlation matrixes were visualized within the R package “corrplot” (Wei et al., 2017). Only correlations were plotted with a P-value of <0.05 and coefficient values R ≤ −0.5 and ≥0.5. Furthermore, M2IA online platform ^92^ was used for the global similarity analyses (PA plot) between metabolome and microbiota data.

#### Transfer experiments

Mice were gavaged with cecal microbiota at ZT13 from either cecal content (CT/ZT13) of *Bmal1^IEC-/-^* mice (n=4, mixture), or control mice kept in LD (n=5, mixture), DD (n=4) or starvation (n=5) into germ-free wild type C57BL6 recipient mice. 100μl of 7×10^6^ bacteria/μl were used for gavaging each mouse. Mice were weekly monitored for bodyweight changes and feces was sampled at week 5 after gavage in DNA stabilizer. Afterwards, mice were sacrificed at the 2^nd^ day in DD at CT13. Native fecal samples for targeted metabolomics were taken at ZT4 and ZT16.

#### Immune cell isolation from the lamina propria

Immune cells were isolated from freshly isolated jejunum and colon. Intestinal tissues were flipped, washed out and cut into 1 cm pieces. To remove epithelial cells, pieces were incubated in DMEM with 20 μL of 1M DTT. After shaking for 15 minutes, tissues were incubated PBS with 200 μL of 150mM EDTA at 37°C with shaking. After 3 times harsh shaking in Hanks buffer, jejunum tissue was then digested at 37°C for approximately 15 min in Thermoshake (200rpm) with 0.6 mg/ml type VIII collagenase (Sigma-Aldrich). Colonic tissue was simultaneously digested under the same condition for approximately 25 min but with the mixture of 0.85 mg/ml type V collagenase (Sigma-Aldrich), 1.25 mg/ml collagenase D (Sigma-Aldrich), 10μl/ml Amphotericin (100x) 1 mg/mL Dispase II, and 30U/ml Deoxyribonuclease I (Sigma). Following digestion, intestinal cells were passed through a 40 μm strainer. Consequently, cells were fixed with 2% PFA, washed, and stored in RPMI at 4 °C until further processing.

#### Fluorescence-activated cell sorting (FACS)

For intracellular stainings, cells were permeabilized with saponin 0.5% and stained with anti CD8 PE-, anti CD3 PerCP/Cy5.5, anti CD4 FITC-, anti IL-17a PE/cy7-, anti INFy-APC conjugated antibodies at dilution 1/100-1/50 for 30 min.

Surface stainings were performed using anti CD11c PE-, anti CD11b APC Cy7-, anti F4/80 PE/cy7-, anti Ly6G APC conjugated antibodies at dilution 1/50 for 30 minutes. Cells were analyzed on a LSR-II (BD Biosciences) flow cytometer and analysis was performed using FlowJo software (FlowJo, LLC).

#### Histology

In formalin fixated tissues in paraffine were cut into 5 μm thick slices and consequently stained according to the following steps: xylene/ 5 min, xylene/ 5 min, Ethanol 100%/ 5 min, Ethanol 100%/ 5 min, Ethanol 96%/ 2 min, Ethanol 96%/ 2 min, Ethanol 70%/ 2 min, Ethanol 70%/ 2 min, Water/ 30 s, hematoxylin/ 2 min, tap water/ 15 s, Scotts Tap Water/ 30 s, Water/ 30 s, Ethanol 96%/ 30 s, Eosin/ 30 s, Ethanol 96%/ 30s, Ethanol 96%/ 30 s, Ethanol 100%/ 30 s, Ethanol 100%/ 30 s, Xylene/ 90 s, Xylene/ 90s (Leica ST5020 multistainer). DPX new mounting media (Merck) was added to preserve the tissues. Stained slides were scanned and further analyzed for histological scoring. Histological scores were assessed blindly based on the degree of immune cell infiltration of all colonic wall layers (mucosa, submucosa and muscularis), crypt hyperplasia, goblet cell depletion and mucosal damage, resulting in a score from 0 (not inflamed) to 12 (severely inflamed) according to Katakura method ^93^.

#### Immunofluorescence staining of CD3

Antigens retrieval of the de-paraffinized and re-hydrated sections was performed by heating the slices in citrate buffer for 34 minutes. After multiple washing steps and tissues were blocked by donkey blocking buffer (1h at room temperature and 1hour at 4C). Tissue sections were incubated overnight with Anti E-cadherin (1:300, mouse, Abcam) anti-CD3 (1:400, rabbit, sigma). Tissue were washed with PBS and followed by incubation with secondary antibodies (donkey anti rabbit 546, donkey anti mouse 647, both from invitrogen with a dilution 1:200) for 1h at room temperature. Finally, DAPI (Sigma) were used to stain the nuclei and then tissues were mounted using Aquatex. Sections were visualized with Fluoview FV10i microscope (Olympus, Shinjuku, Japan).

#### PICRUST 2.0

For prediction of functional of metagenomic functionality. Sequence of the gut controlled zOTUs, captured as described above, were used to construct the metagenome using PICRUST2.0 ^94^. Corrected zOTU 16s rRNA gene copy number is multiplied by the predicted functionality to predicted the metagenome. Resulted enzymatic genes classified according to Enzyme Commission (EC) numbers were mapped to Metacyc pathways. Superclasses were removed and Metacyc pathways abundance was used for statistical analysis using STAMP (2.1.3). Statistical differences were calculated based on White’s non-parametric t-test and Benjamini Hochberg dales discovery rate to adjusted for multiple testing.

### Statistical Analyses

Statistical analyses were preformed using GraphPad Prism, version 9.0.0 (GraphPad Software), JTK_cycle v3.1.R (^25^) or R. With the use of the pipeline Rhea (Lagkouvardos) between-sample microbiota diversity is calculated by generalized UniFrac using GUniFrac v1.1. distances. Quantification of GUniFrac distances was always done in comparison to ZT1 or CT1. Circadian and diurnal patterns of individual 24h period graphs as well as phase calculation were analysed by fitting a cosine-wave equation: y=baseline+(amplitude·cos(2?π?((x-[phase shift)/24))), with a fixed 24-h period for phyla, richness and single zOTU or by using the non-parametric algorithm JTK_cycle for overall rhythmicity of all zOTUs. Connected straight lines in of individual 24h period graphs within the figures indicate significant rhythmicity based on cosine analyses whereas dashed lines indicate non-significant cosine fit. Comparison of rhythm between data sets were preformed using an adjusted version of the Compare rhythm script ^26^, which performs JTK_cycle followed by DODR ^95^. Some CCGs when indicated were analysed with a harmonic - regression: y= baseline + (amplitude A * cos (2·π· ((x - [phaseshift A) / 24))) + (amplitude B * cos (4·π·((x - [phase shift B) / 24))). Amplitude calculations depicted in the manhattan plots are based on the output of JTK_cycle and the phase was calculated by cosine-wave regression. Analysis between two groups was performed using the non-parametric Mann-Whitney test. A p value ≤0.05 was assumed as statistically significant.

Heatmaps were generated using the online tool “heatmapper.ca”^96^. Heatmaps were sorted based on peak phase of controls. Abundance plots were generated using SIAMCAT package in R using the “check.associations() function”^97^. Visualisation of zOTUs and samples trees were conducted using the online platform “evolgenius.info”^89^.

## Data availability

Microbiota sequencing data and metabolite data will be available from the Sequence Read Archive (SRA) and the MetaboLights database for Metabolomics experiments (https://www.ebi.ac.uk/metabolights/) upon request. The modified version of the compare rhythm script is availabe (https://github.com) upon request.

