## Supplemental Figure 1-4 for "The intestinal circadian clock drives microbial rhythmicity to maintain gastrointestinal homeostasis"

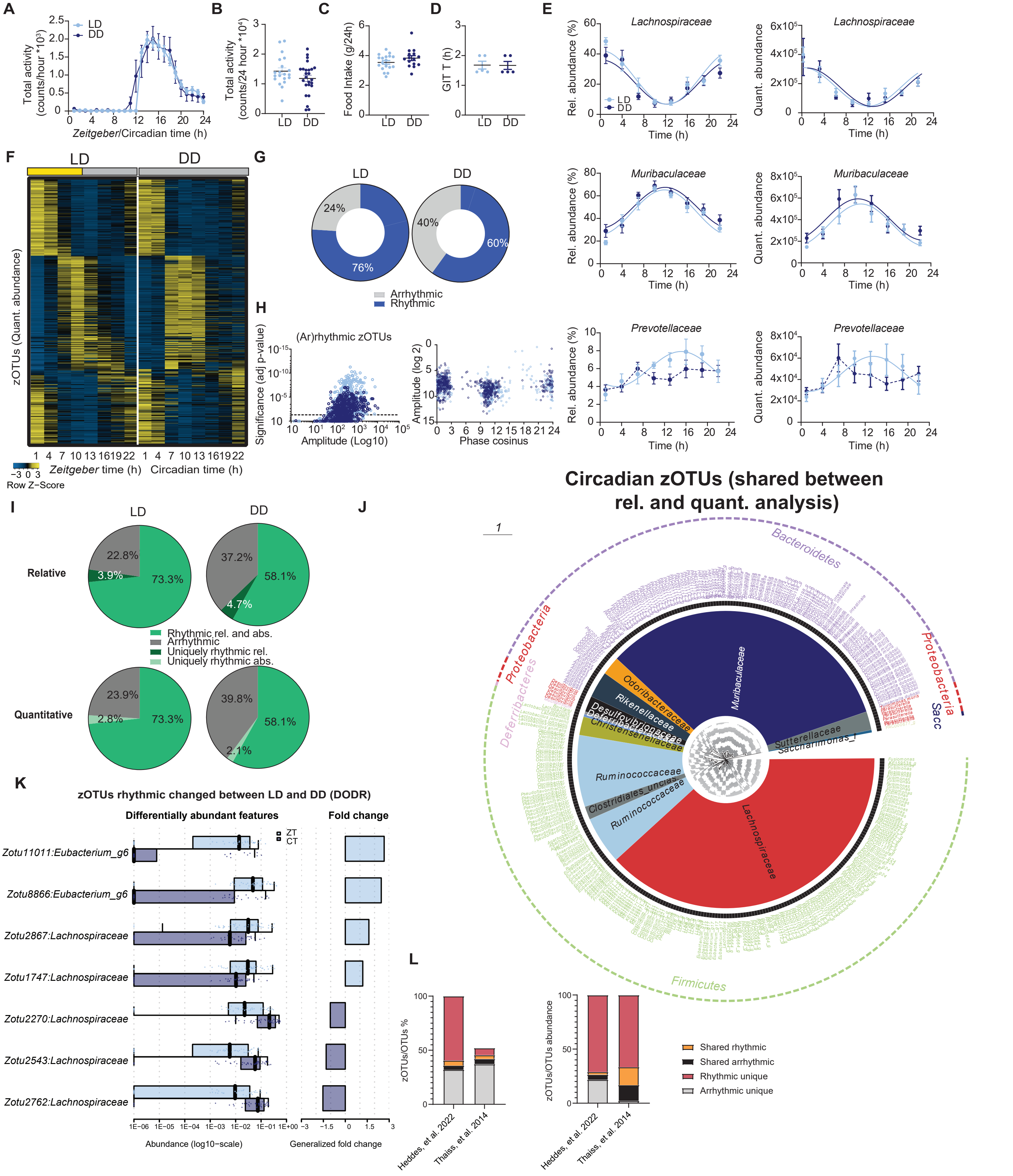

**Supplement Figure 1 Diurnal and circadian rhythms in behavior and microbiota composition** (A) Diurnal (LD) and circadian (DD) total wheel-running activity profiles and 24-h summary (B) (n = 20-25/condition). Total daily food intake (n = 15-17/condition) (C) and gastro-intestinal transit time (GITT) (n = 5/condition) (D). (E) Diurnal and circadian profile of relative (left) and quantitative (right) abundance of the major family of fecal microbiota. (F) Heatmap depicting the quantitative abundance of 580 zOTUs (mean relative abundance > 0.1%; prevalence > 10%). Data are normalized to the peak of each zOTU and ordered by the peak phase in LD conditions. (G) Pie-charts indicate the amount of rhythmic (blue) and arrhythmic (grey) zOTUs, identified as rhythmic (adj. p-value ≤ 0.05) by JTK_Cycle. (H) Significance and amplitude of rhythmic and arrhythmic zOTUs (left) and phase distribution (right) in LD and DD based on quantitative analysis. Dashed line indicates adj. p-value = 0.05 based on JTK_Cycle. (I) Pie charts indicating percentage of overlap in rhythmic (green) and arrhythmic (grey) zOTUs between relative (top) and quantitative (bottom) analyses in LD (left) and DD (right) conditions, identified as rhythmic (adj. p-value ≤ 0.05) by JTK_Cycle. (J) Taxonomic tree of circadian zOTUs shared by both relative and quantitative analyses. Taxonomic ranks are from phylum (outer dashed ring), family (inner ring) to genera (middle, color coded according to phylum) indicated by individual branches. (K) Box and bar plots illustrate the alteration in relative abundance and fold change between LD and DD of zOTUs (Wilcoxon, adj. p-value ≤ 0.05), which showed altered rhythmicity according to the adjusted compare rhythm script based on DODR. (L) Bar charts comparing rhythmic/arrhythmic zOTUs/OTUs percentage (left) and abundance (right) of Thaiss et al. and Heddes et al. Data were normalized to the amount of zOTUs for percentage calculation, identified as rhythmic (adj. p-value ≤ 0.05) by JTK_Cycle. Significant rhythms are illustrated with fitted cosine-regression solid line; data points connected by dotted lines indicate no significant cosine fit curve (p-value > 0.05) and thus no rhythmicity. LD (light-blue) and DD (dark-blue). n = 6 mice/time point/light condition unless otherwise indicated. Data are represented as mean ± SEM.

### Supplemental Figure 2

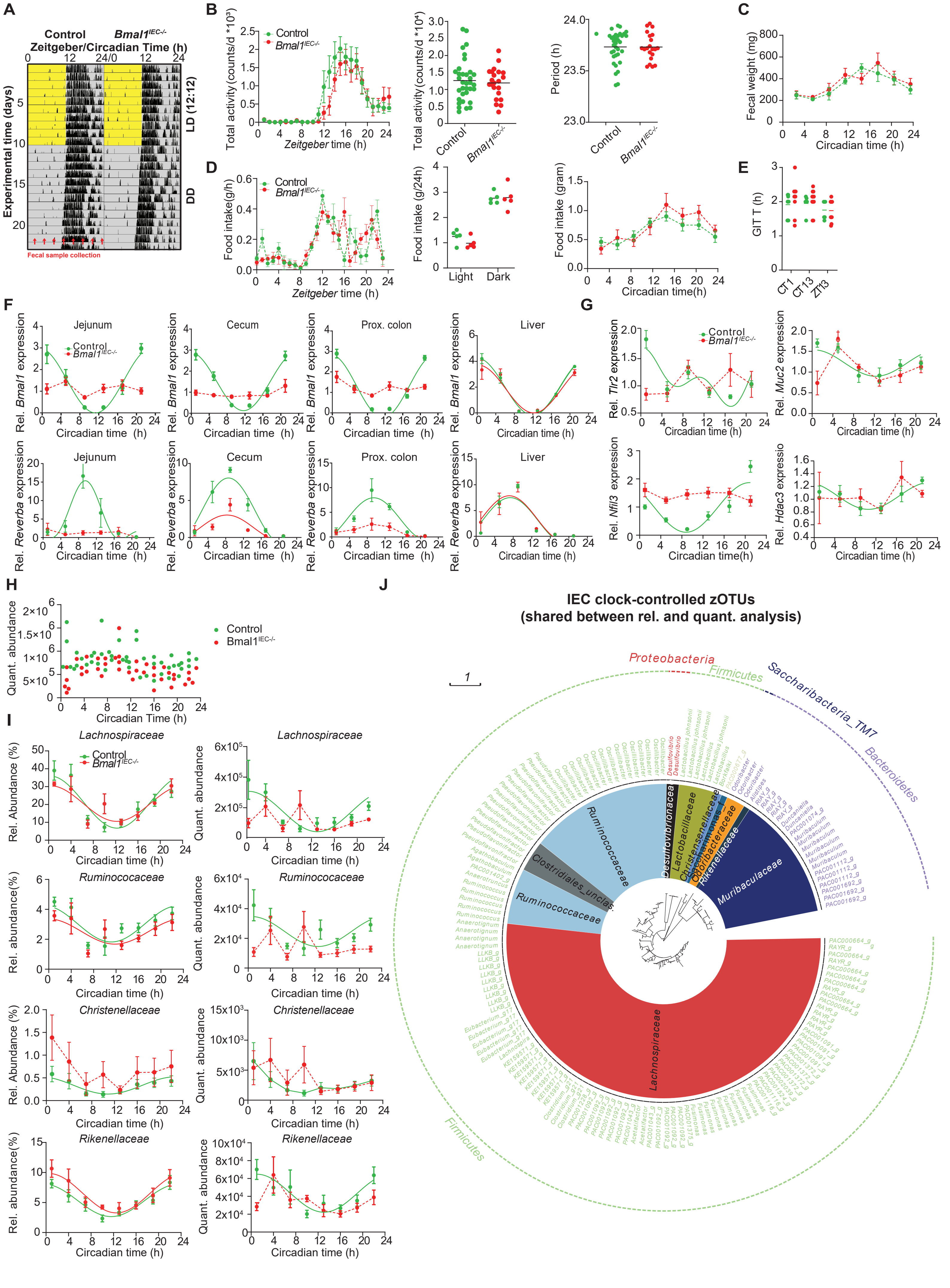

**Supplement Figure 2 Characterization of rhythmic behavior and microbial profiling of *Bmal1$^{IEC-/-}$* mice** (A) Representative actogram in LD and DD conditions of *Bmal1$^{IECfl/fl}$* controls and *Bmal1$^{IEC-/-}$* mice, red arrows indicate fecal samples collection time points. (B) Activity profile of *Bmal1$^{IEC-/-}$* (n = 22) and control mice (n = 33) in light-dark (LD) cycle (left) and the quantification of circadian activity (middle) and the period in DD (right) as well as (C) fecal weight in DD over time and (D) food intake diurnal profile and its average (middle), and food intake circadian profile (right) (n = 5/genotype). (E) GITT in LD and DD (n = 6-8). Expression profiles of core clock genes (F) and clock-controlled genes (G). (H) 16S copy number over time (2-way ANOVA) (I) Circadian profiles at family level of relative abundance (left) and quantitative abundance (right) of control and *Bmal1$^{IEC-/-}$* mice. (J) Taxonomic tree of fecal circadian gut clock controlled microbiota uniquely rhythmic in control mice in both relative and quantitative analyses. Taxonomic ranks are from phylum (outer dashed ring), family (inner ring highlighted) to genera (middle, color coded according to phylum) which are indicated by the individual branches. Significant rhythms are illustrated with fitted cosine-regression or fitted harmonic-regression; data points connected by dotted lines indicate no significant cosine fit curves (p-value > 0.05) and thus no rhythmicity. *Bmal1$^{IEC-/-}$* (red) and control (green). n = 5-6 mice/time point/genotype. Data are represented as mean ± SEM. Significance: p-value ≤ 0.05.

### Supplemantary Figure 3

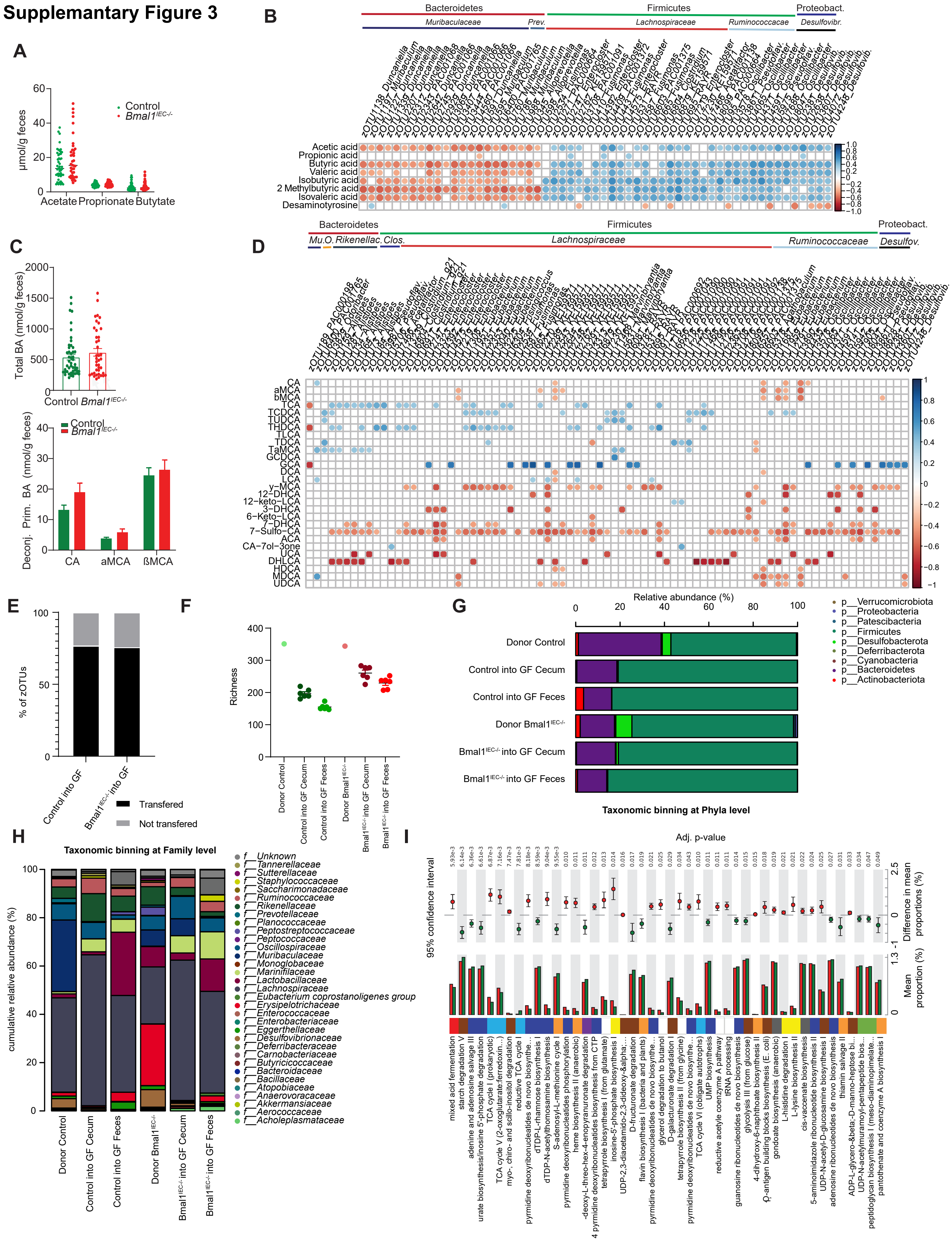

**Supplement Figure 3. Arrhythmic microbial transfer to germ-free mice.** (A) Total SCFA concentrations in feces. (B) Spearman correlation of SCFA (p-value ≤ 0.05 and R ≤ -0.5 (red) or R ≥ 0.5 (blue)) with gut-controlled bacteria taxa. (C) total and deconjugated bile-acid levels in feces and their (D) Spearman correlation of bile-acids (p-value ≤ 0.05 and R ≤ -0.5 (red) or R ≥ 0.5 (blue)) with gut-controlled bacteria taxa. n = 5-6 mice/time point/genotype (repeated measures). Control (green) and *Bmal1*$^{IEC-/-}$ (red).(E) Percentage of zOTUs transferred (grey) into recipient mice (n = 6/genotype). (F) Richness of donor (n = 4 mixture) and recipient samples collected at CT13/ZT13. Taxonomic binding of microbiota from donor and recipient mice at CT13/ZT13 at phyla (G) and family (H) level. (I) Pathways predicted using PICRUST2.0 on intestine clock-controlled zOTUs showing significant differences in abundance between genotypes. Pathways are colored according to their sub-class. Statistical differences for Picrust data were calculated based on White's non-parametric t-test and Benjamini Hochberg dales discovery rate to adjusted for multiple testing. Cholic acid (CA), a-Muricholic acid (aMCA), b-Muricholic acid (bMCA), Taurocholic acid (TCA), Taurochenodeoxycholic acid (TCDCA), Tauroursodeoxycholic acid (TUDCA), Taurohyodeoxycholic acid (THDCA), Taurolithocholic acid (TLCA), Taurodeoxycholic acid (TDCA), Tauro-a-Muricholic acid (TaMCA), Glycochenodeoxycholic acid (GCDCA), Glycocholic acid (GCA), Deoxycholic acid (DCA), Lithocholic acid (LCA), y-Muricholic acid (y-MCA), 12-Dehydrocholic acid (12-DHCA), 12-Ketolithocholic acid (12-keto-LCA), 3-Dehydrocholic acid (3-DHCA), 6-Ketolithocholic acid (6-keto-LCA), 7-Dehydrocholic acid (7-DHCA), 7-Sulfocholic acid (7-sulfo-CA), Allocholic acid (ACA), Cholic-acid-7ol-3one (CA-7ol-3one), Ursocholic acid (UCA), Dehydrolithocholic acid (DHLCA), Hyodeoxycholic acid (HDCA), Murideoxycholic acid (MDCA), Ursodeoxycholic acid (UDCA).

### Supplement Figure 4

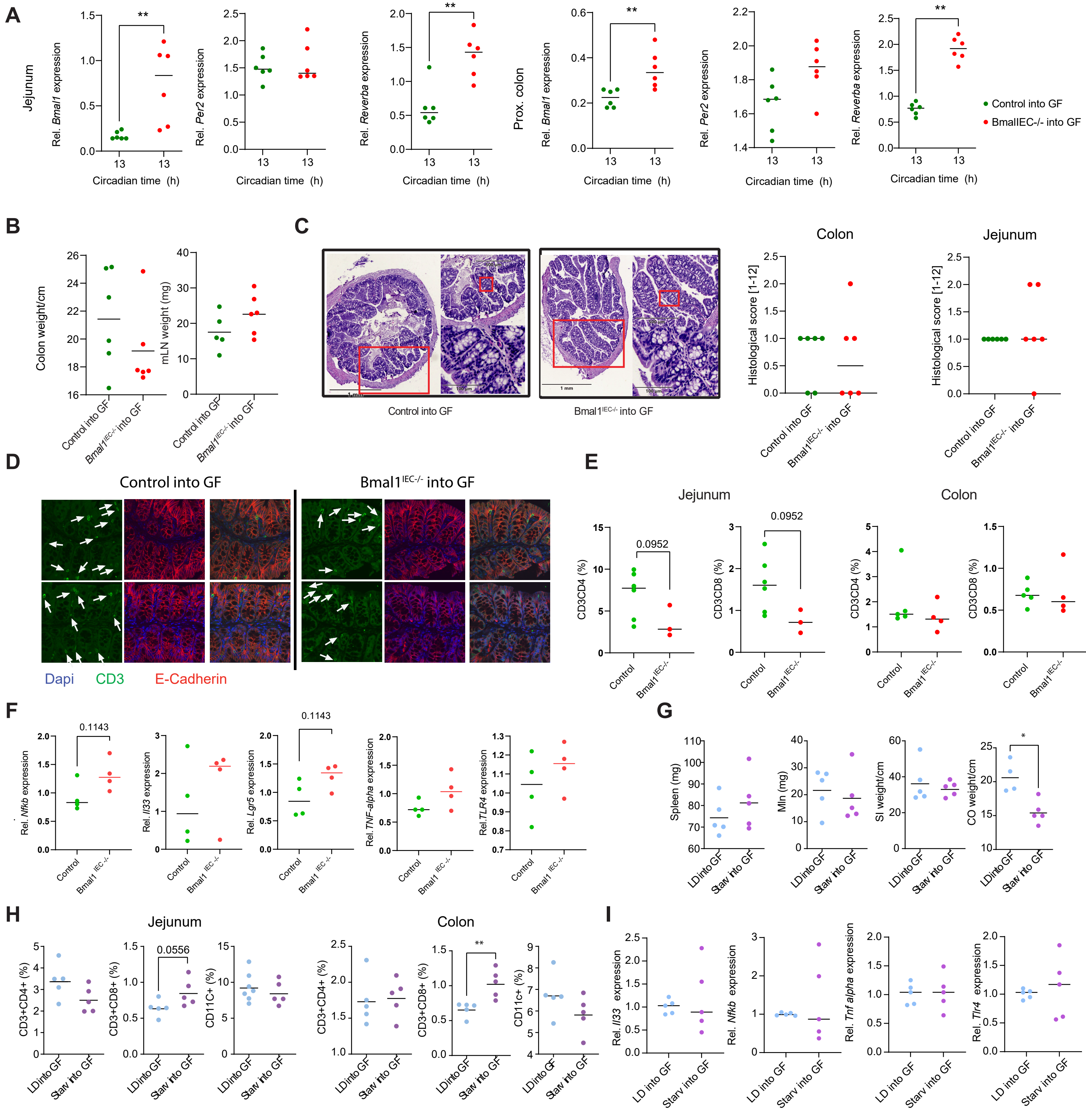


**Supplement Figure 4. Immune cell recruitment and gene expression in SPF donors and germ free mice after microbiota transfer** (A) Clock gene expression measured at CT13 in Jejunum (left) and proximal colon (right) of recipient mice 6 weeks after microbiota transfer of *Bmal1*$^{IECfl/fl}$ control or *Bmal1*$^{IEC-/-}$ mice (n = 6/genotype). (B) Organ weights of recipient mice after receiving control or *Bmal1*$^{IEC-/-}$ cecal microbiota (n = 6/genotype). (C) Cross section of proximal colon along with the histological scoring of proximal colon and jejunum of germ-free mice after receiving control or *Bmal1*$^{IEC-/-}$ cecal microbiota (n = 6/genotype). (D) Immunofluorescence staining of CD3 (green), Ecadherin (red) and Dapi (blue) of proximal colon of germ-free mice after receiving control or *Bmal1*$^{IEC-/-}$ cecal microbiota. (E) Frequency of CD3+CD4+, CD3+CD8+ cells in jejunum and colon of SPF (donor) *Bmal1*$^{IEC-/-}$ and control mice (n = 3-4/genotype). (F) Relative gene expression of Tlr4, Tnfa, Il33, Nfkb, Lgr5 in the proximal colon of SPF control and *Bmal1*$^{IEC-/-}$ mice (n = 4/genotype).(H) Frequency of CD3+CD4+, CD3+CD8+ and CD11c+ cells in jejunum and colon after transfer of LD microbiota and starvation microbiota into GF-BL6 recipients (n = 5/genotype). (I) Organ weights of recipient mice after receiving LD or starvation microbiota (n = 5/genotype). (I) Relative gene expression of Tlr4, Tnfa, Il33, Nfkb, Ang4 in the proximal colon into LD microbiota and starvation microbiota recipient mice (n = 5/genotype). Data are represented as mean ± SEM. * p ≤ 0.05, ** p ≤ 0.01 (Mann-Whitney U test).
