## Supplementary figures and images for "The intestinal circadian clock drives microbial rhythmicity to maintain gastrointestinal homeostasis"

### Supplemental Information Figure 1

Supplementary Information Fig. 1

A

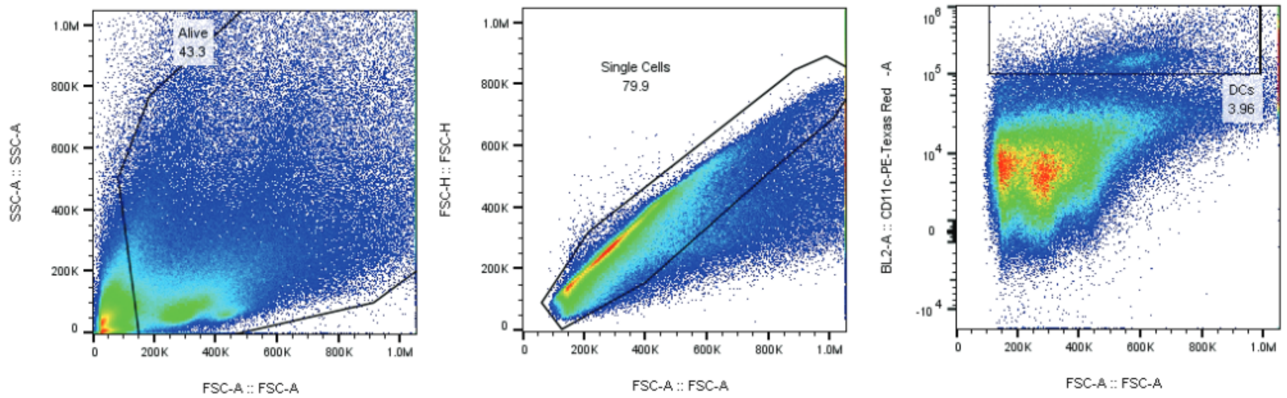


B

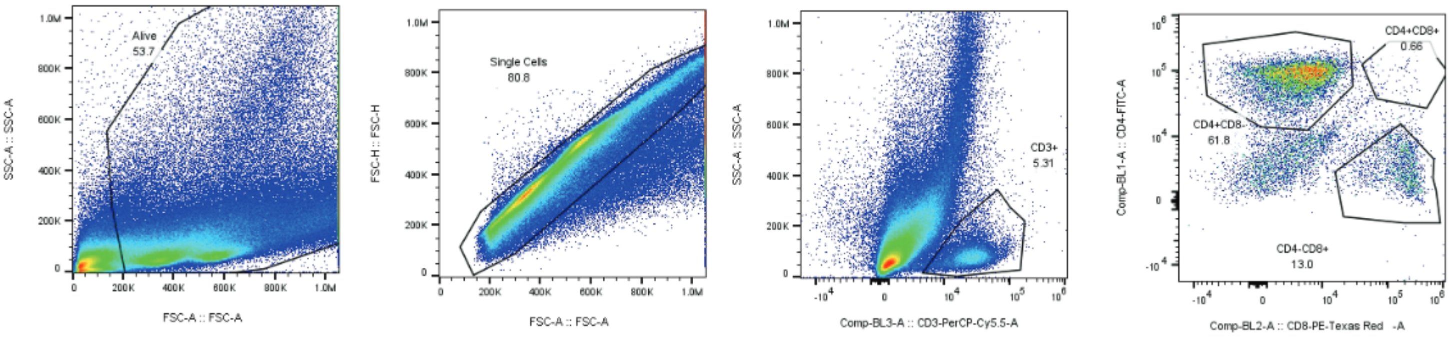
